# Loss of myosin VI leads to age-dependent changes in the maintenance of heart ventricle mitochondria

**DOI:** 10.64898/2026.10.13.758738

**Authors:** Olena Karatsai-Miaskowska, Lilya Lehka, Jolanta Nowak, Weronika Ficerman-Kuta, Dominika Wojton, Krzysztof Skowronek, Adam Szewczyk, Maria Jolanta Redowicz

## Abstract

**Aims:** We have recently shown that unconventional myosin VI (MVI), a unique actin-based molecular motor, is involved in age-progressive cardiac dysfunction. Here, we aimed at elucidation of mechanisms of MVI involvement in this phenotype. We propose that MVI plays a role as a regulator of mitochondria quality control as well as of their organization and distribution in the heart, an organ with extremely high energy demand.

**Methods and results:** We analyzed heart samples from MVI knockout mice (Snell’s waltzer) and their heterozygous littermates: newborns (P0), 3-month-old (3m), and 12-month-old (12m) mice. Transmission electron microscopy and immunostaining revealed that loss of MVI affected mitochondria network organization and promoted mitochondria clustering in the perinuclear space. Moreover, accumulation of large functional mitochondria, increase in total mitochondria mass with a concomitant increase in the amount of damaged mitochondria were observed in 12m hearts. The Western blot technique revealed that loss of MVI, at all examined time points, did not affect the activity of AMPK, the key energy balance sensor, and the level of the respiratory chain components. However, in P0 samples it affected the levels of OPA1, involved in mitochondria fusion, and the activity of S6, the downstream effector of mTOR. Also, loss of MVI affected autophagy/mitophagy activity in P0 hearts, leading to accumulation of autophagosomes. A pull-down assay enabled identification of over three hundred potential MVI-binding partners; among them were proteins involved in mitochondria organization and functions, autophagy, ion transport, and cell respiration.

**Conclusion:** Taken together, our results indicate that loss of MVI evokes changes disclosed already in newborns that lead to profound alterations in the mitochondria organization and maintenance with age. We believe that accumulation of changes in mitochondria structure, distribution and quality commits to the age-progressing cardiac remodeling and dysfunction.

**Graphical Abstract:** 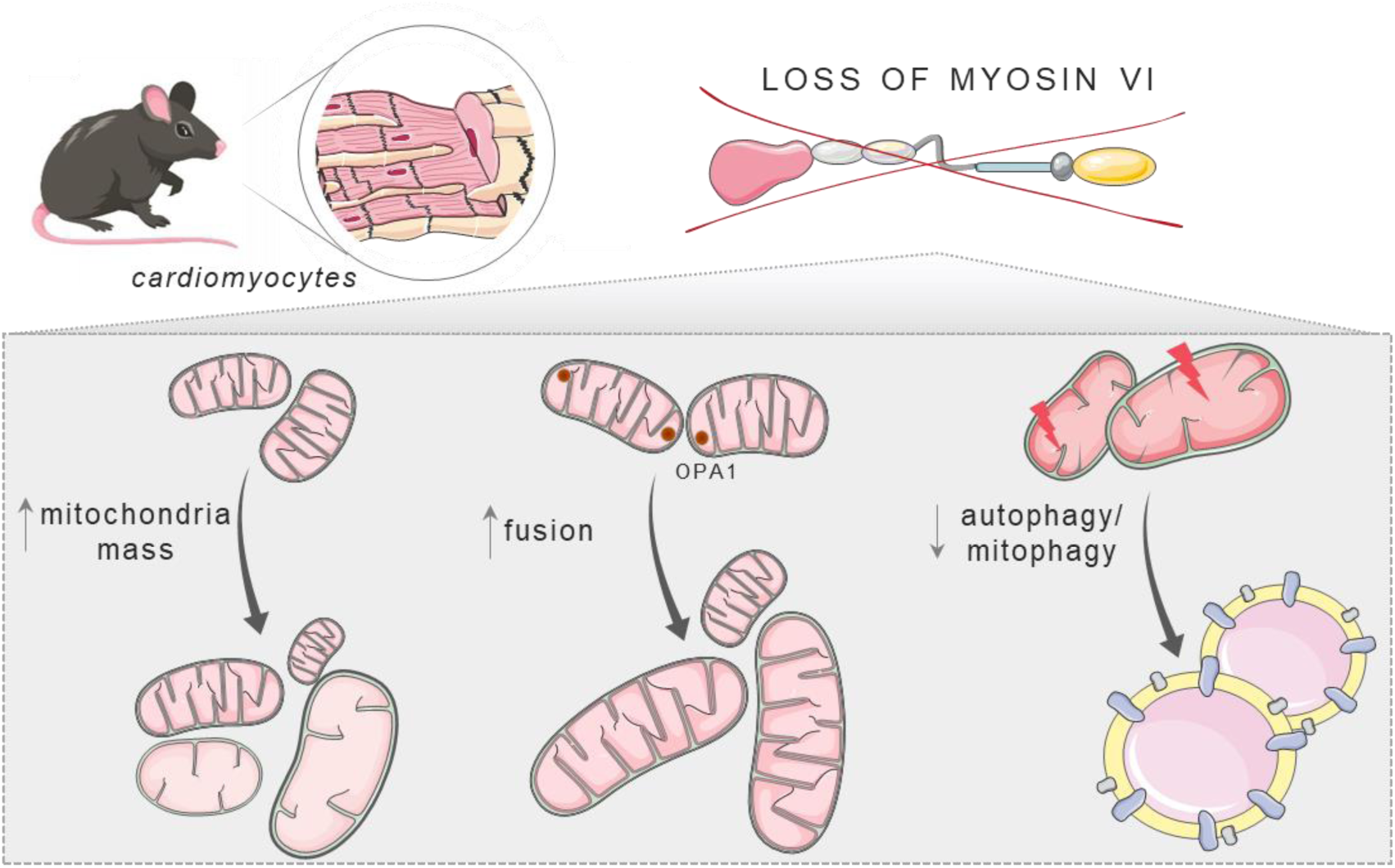

## 1. Introduction

Proper heart functioning is highly dependent on the continuous energy production. Mitochondria are considered as the “powerhouses” of eukaryotic cells and, therefore, the key element in the regulation of cell and tissue functioning, including the heart, an organ with very high energy demand. In adult cardiomyocytes mitochondria occupy more than 30% of the cell volume and produce ATP through oxidative phosphorylation (OXPHOS) ^1^. Mitochondria are also important calcium-buffering organelles, what makes them the key regulator of cardiac function through modulation of myocardium contraction, nutrient flux, regulation of cell cycle, apoptosis, and development ^1,2^. In adult cardiomyocytes, distribution and organization of mitochondria are essential to maintain cellular functions whereas distinct mitochondrial subpopulations are differently involved in the cellular processes ^3^.

Mitochondria functioning has to be precisely controlled to avoid cell/organ impairment. Multiple mechanisms are involved to maintain mitochondria homeostasis such as the system of quality control (mitochondria biogenesis, fusion/fission events, and mitophagy), changes in mitochondria metabolism, interaction network with other cellular compartments as well as communication with the nucleus ^4^. In many cell types, mitochondria are highly dynamic organelles as fusion and fission events continuously change their network and interconnection with other organelles modulating cellular functions ^4,5^. In cardiomyocytes, the intensity of mitochondrial dynamics and distribution are dependent on the stage of cardiomyocyte maturation and proteins involved in mitochondrial dynamics are also important in the maintaining of other vital functions, such as mitochondrial biogenesis, organization of the inner membrane, and apoptosis ^4,6^. Mitophagy, a specialized type of autophagy, in turn, provides a selective targeting and elimination of dysfunctional or damaged mitochondria ^7,8^. This process is a core of mitochondrial integrity and cellular homeostasis maintenance, playing a pivotal role in cardiomyocytes ^7^.

A large number of diverse proteins maintains normal mitochondria structure and function. It has been shown that any disturbance in their functioning leads to arrhythmia, heart failure, ischemic heart disease, hypertension, and the development of cardiomyopathies ^1,7,9^. Also, any impairment of mitochondria as well related to endothelial dysfunction could result in atherosclerosis ^7^. Both the increase and accumulation of dysfunctional mitochondria are the basis for various cardiovascular diseases, including cardiomyopathy ^7,10^.

Unconventional myosin VI (MVI) is a unique actin-interacting molecular motor moving toward the minus end of actin filaments ^8^. MVI contains N-terminal motor domain, a neck with two calmodulin (CaM)-binding sites, and C-terminal tail which includes 3-helix bundle, single α helix, distal tail region, and the cargo-binding domain (*Figure 1A*). Organization of the tail domain plays a crucial role in MVI interactions with specific cargos (binding partners) within the cell, and thus enabling MVI to be involved in a panoply of vital processes ^11–14^.

**Figure 1.**
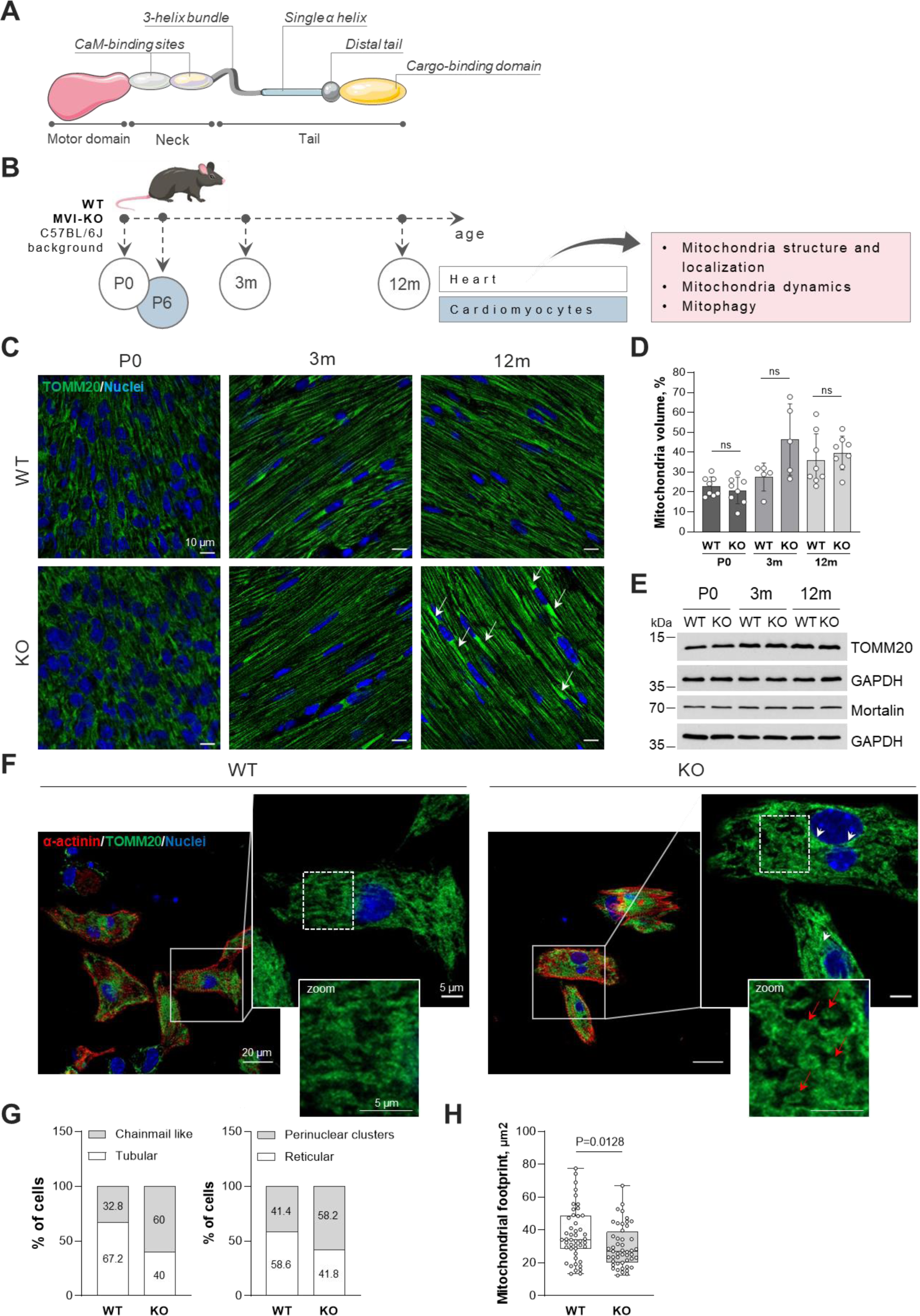
Effect of MVI loss on the mitochondria network in murine cardiomyocytes. **A**, schematic structure of the myosin VI (MVI) monomer; detailed explanation in the text. **B**, schematic presentation of experimental design. Heart ventricles (hearts) were isolated from Snell’s Waltzer mice (C57BL/6J background) not synthesizing functional MVI (MVI-KO) with respect to control (WT), heterozygous littermates at different age: newborns (P0, both sexes), 3-month-(3m) and 12-month-old (12m) male mice. 6-days-old mice of both sexes were used for cardiomyocyte isolation. Evaluation of mitochondria structure and localization, dynamics, and mitophagy was performed. **C**, representative immunofluorescence staining for the mitochondria marker, TOMM20 of longitudinally oriented cardiomyocytes within transverse heart sections. Nuclei were labeled with DAPI. *White arrows* show clustered mitochondria in the perinuclear space. **D**, quantification of panel **C**, showing a mitochondria volume (TOMM20 stained area) in cardiomyocytes. Results were expressed as means ±SD. Each circle (°) represents an independent experiment (N ≥ 5). Statistical significance was analyzed with parametric or non-parametric t test. ns – not statistically significant difference. **E**, representative Western blots of the mitochondria markers. GAPDH was used as an internal loading control. **F**, double staining for cardiomyocyte marker, α-actinin and TOMM20 of primary cultured cardiomyocytes. *White arrows* show clustered mitochondria in the perinuclear space. *Red arrows* show chainmail like mitochondrial morphology in MVI-KO cardiomyocytes. *Dashed square* shows a magnified area. **G**, a number of cells (in % of total amount) with chainmail like mitochondrial morphology (left graph) and with mitochondrial clusters within perinuclear space (right graph). **H**, mitochondria footprint was quantified using Mitochondria Network Analysis (MiNA) tool. Results are shown as box and whisker plots, horizontal line indicates median. Each circle (°) represents each analyzed cell. Statistical significance was analyzed with the use of Mann-Whitney test. P value is provided on the graph. In **G** and **H,** the analysis was performed on 62 WT and 55 MVI-KO cells from 5 independent experiments.

Snell’s waltzer (SV) mice (C57BL/6J background) are natural MVI-knockout (MVI-KO) animal model exhibiting head-tossing, circling, and hyperactivity ^15^. Recently, we showed that MVI is an important player in heart functioning, heart growth, and development. We showed that lack of MVI leads to the age-dependent progressive heart remodeling: from mild hypertrophy to dilated-like cardiomyopathy. These changes were supported by an increase in cell proliferation in newborns, impairment of intercalated disc integrity, and systolic dysfunction (Karatsai et al. 2023). Furthermore, MVI is associated with the maintenance of mitochondria biogenesis and energy production in skeletal muscles through AMPK signaling, and is involved in the selective transport of damaged mitochondria in fibroblasts ^8,12,16^.

Taken together, the observations described above suggest the existence of an important link between MVI and mitochondria homeostasis. In this study, we describe that loss of MVI in the heart affects mitochondria organization and distribution as well as mechanisms of mitochondria quality control, such as fission/fusion events and mitophagy without a major effect on the global energy balance. Moreover, we show that the changes in mitochondria accumulation and distribution are progressing with age.

## 2. Methods

### 2.1 Animals

All experiments were performed on Snell’s waltzer (SV) mice (C57BL/6J background) not synthesizing functional myosin VI (MVI) with respect to a control, heterozygous littermates of different ages: newborns (P0, mice of both sexes), 3-month-(3m, males), and 12-month-old (12m, males) mice. 6-days-old (P6, both sexes) mice were used for cardiomyocytes isolation. SV mice serve as a natural knockout model for MVI. The spontaneous mutation introducing a premature stop codon in the cargo-binding tail leading to a lack of functional MVI ^17^. Heart ventricles (next hearts) were isolated, washed in ice-cold PBS, gently drained, immediately frozen on the dry ice, and stored at −80°C until used. Animal housing and sacrifice procedures were performed in compliance with the European Communities Council directives adopted by the Polish Parliament (Act of 15 January 2015 on the protection of animals used for scientific or educational purposes (consolidated text Journal of Laws 2021 item 1331, as amended)). Animals were living on the standard rodent chow diet and housed under pathogen-free conditions in the animal facility of the Nencki Institute. Mice were euthanized by inhalation of isoflurane (Iso-Vet 1000 mg/g; Piramal Critical Care) at a concentration of 5% until deep anesthesia and unresponsiveness were achived, followed by cervical dislocation as a secondary method for euthanasia. Isoflurane was administrated once for euthanasia. The following internal approvals were granted by the Director of the Nencki Institute of Experimental Biology: 88W/2021/IBD (O.K.M.), 410W/2021/IBD (L.L.), 530W/2024/IBD (W.F.K.), and 155/2025/IBD (M.J.R). We have made all efforts to minimize the suffering of the animals and used the smallest number of animals to obtain reliable results.

### 2.2 Genotyping

Tail clips obtained from P0 mice were used for genotyping according to manufacturer’s instructions (Selleckchem, B40013). Primers for MVI genomic DNA amplification are listed in Supplementary material online, *Table S1.* The PCR mixture contained 1 µl of genomic DNA, 7.5 µl of ddH_2_O, 10 µl of 2 x M-PCR OPTI^TM^ Mix 0.5 µM SV1, SV2, and SV3 primers. PCR was performed as follows: 5 min of activation at 94°C followed by 35 cycles of 20-s denaturation at 94°C, 30 s of annealing at 56°C, 30 s of extension at 72°C, and one 5-min finish cycle at 72°C. After the PCR reaction, DNA was subjected to electrophoretic separation in a 2% agarose. Mice with *^sv/sv^* genotype (homozygous, MVI-KO) displayed a 318 bp genomic product, while animals with *^+/sv^*genotype (heterozygous, WT) displayed two genomic products: 318 and 230 bp.

### 2.3 Cell culture

Murine cardiomyocytes were isolated from 6-days-old (P6) Snell’s waltzer (SV) mice. Hearts were separated, washed with PBS (VWR, 392-0434) supplemented with 1% penicillin/streptomycin (P/S; Gibco, 15140-122) and 20 mM 2,3-butanedione 2-monoxime (BDM; Sigma-Aldrich, B0753) to increase survivability of cardiomyocytes, and then transferred to the PBS+P/S+BDM mixture with few drops of betadine (10% Polyvidonum iodinatum solution). Next, hearts were cut into small pieces and transferred to 0.25% trypsin-EDTA (Gibco, 25200-056) with 1% P/S and 20 mM BDM for overnight incubation at 4°C. Then, trypsin solution was carefully aspirated and collagenase type II (0.75 mg/ml, 1 ml/heart; Gibco, 17101-015) supplemented with DNAse I (10 µg/ml; Roche, 11284932001) was added, followed by a gentle pipetting and incubation at 37°C for 5 min. Upper part of suspension was transferred to Plating Medium (DMEM high glucose supplemented with GlutaMAX-1 (Gibco, 31966-021), 4% FBS (Gibco, 10084-147), and 1% P/S). If necessary, more additional digestion steps were carried out. Cell suspension was then passed through the 100-µm cell strainer (VWR, 76327-100) and centrifuged two times at 200 × g for 8 min at 21°C. Pellet was resuspended in Plating Medium. The single cells suspension was plated into not covered dish for 3 h at 37°C with 5% CO_2_ to get rid of fibroblasts. After that, unattached cells (cardiomyocytes) were collected, counted, and seeded into 0.02% gelatin/2 µg/ml fibronectin (EmbryoMax® Gelatin Solution; Millipore, ES-006-B/Bovine Fibronectin; Sigma-Aldrich, F1141) coated glass coverslips (VWR, 631-0153) in 12-well plate at a density of 450 × 10^3^ cells/well. The homogeneity of the cardiomyocyte population was evaluated on day 2-3 by immunofluorescence staining for the cardiomyocyte marker, α-actinin, and fibroblast marker, vimentin (see Supplementary material online, *Table S2 and S3*). The analysis revealed that around 40% of cells were cardiomyocytes (Supplementary material online, *Figure S1*).

### 2.4 Transmission electron microscopy

Hearts were isolated and washed in ice-cold PBS in order to get rid of the blood contamination. Then hearts were gently drained and immediately placed in fixative solution containing 2.5% glutaraldehyde and 2% paraformaldehyde solution (PFA) in 0.1 M sodium cacodylate buffer for at least 24 h at 4°C. Next, the sample preparation was performed by Dr. Andrzej Szczepankiewicz from the Laboratory of Electron Microscopy, Nencki Institute. Briefly, after fixation, samples were washed three times for 10 min in the cacodylate buffer and cut out from left ventricle at 2 mm x 2 mm thickness. Then, tissue was postfixed in 1% osmium tetroxide for 1 h at room temperature (RT) and stained with 2% uranyl acetate for 40 min. Sample dehydration was performed by incubation in increasing acetone concentrations (25%, 50%, 70%, 90%, 96%, and 2 x 100% for 10 min each). After that, tissue were embedded in the mixture of pure acetone and Epon resin (1:1, 1:2, and 2:1 for 30 min each), and then in pure Epon resin exchanged three times (1 h, overnight, and 1 h). After two days of polymerization at 60°C, transverse cardiac sections were cut at 70 nm thickness on Leica Ultracut R ultramicrotome, collected on formvar TEM 200 mesh copper grids and stained with Reynold’s lead citrate for 2 min.

Micrographs of cardiac tissue were taken using JEOL JEM-1400 (JEOL Co.) Electron Microscope equipped with 11 Megapixel TEM MORADA G2 (EMSIS GmbH). Optical sections (3864 pixels×2688 pixels) were collected using RADIUS 2.2 software (EMSIS GmbH). Images were analyzed using open-source image processing software ImageJ 1.54p (Wayne Rasband and contributors, National Institutes of Health) as described at ^18^. Analysis of mitochondria morphology was performed on low magnification images (8000x) of longitudinally oriented cardiomyocytes. Measurements were repeated in 10 images each from three different individuals.

### 2.5 Immunocytochemical staining

#### 2.5.1 Heart section staining

Heart cryosections were prepared as described earlier ^19^. Before immunostaining, cryosections (10 µm thickness) were air-dried for 30 min at RT and washed with PBS for another 25 min. Next, they were fixed in 4% PFA for 30 min, washed twice with PBS for 15 min, quenched for 30 min with 50 mM NH_4_Cl (in PBS), permeabilized with 0.3% Triton X-100 in PBS for 20 min, and finally incubated for 1.5 h in a blocking buffer (5% horse serum in PBS supplemented with 0.3% Triton X-100 (PBS-T0.3)). After that, samples were incubated overnight at 4°C with appropriate primary antibodies (see Supplementary material online, *Table S2*) diluted in blocking buffer. Next, the sections were washed three times for 15 min with PBS-T0.3 and incubated with corresponding secondary antibodies (see Supplementary material online, *Table S3*) for 1.5 h at RT. Finally, the samples were washed three times for 15 min in PBS-T0.3 and mounted using Vectashield PLUS antifade mounting medium with DAPI (Vector Laboratories, H-2000).

Microscopy analysis was performed using Confocal laser scanning microscope Zeiss LSM780, Inverted Axio Observer Z.1 equipped with Plan Apochromat 63x/1.4 Oil DIC M27 objective. An argon laser 488 nm and diode laser 405 nm (CW/pulsed), were used to excite Alexa Fluor 488 and DAPI fluorescence, respectively. Optical sections (2048 pixels×2048 pixels×16 Bit) were collected at 0.3 µm in z axis. Images were processed using free Zen Lite 3.10 software (Carl Zeiss Microscopy). Mitochondria volume was counted using ImageJ 1.54p software.

#### 2.5.2. Cell staining

Cells cultured on the glass coverslips were gently washed once with warm PBS, fixed with 4% PFA for 20 min, washed twice in PBS, and incubated in 50 mM NH_4_Cl for 30 min. After that, cells were washed once with PBS, permeabilized with 0.2% Triton X-100 in PBS for 10 min, and incubated for 1.5 h in a blocking buffer (2% horse serum in 0.02% Triton X-100 PBS (PBS-T0.02)). Then, samples were incubated with appropriate primary and secondary antibodies (see Supplementary material online, *Table S2 and S3*) accompanied by wash cycles in PBS-T0.02. Finally, samples were mounted by Vectashield medium with DAPI.

Images were collected using Confocal Laser Scanning Microscope with Airyscanning Zeiss LSM800, Inverted Axio Observer.Z1/7 equipped with Plan-Apochromat 40x/1.4 Oil DIC (UV) VIS-IR M27 objective. An argon laser 488 nm, DPSS diode 561 nm, and diode laser 405 nm (CW/pulsed), were used to excite Alexa Fluor 488, 546, and DAPI fluorescence, respectively. Optical sections (1024 pixels×1024 pixels×16 Bit) were collected at 0.3 µm in z axis. Images were processed using free Zen Lite 3.10 software.

Mitochondria morphology was analysed with Mitochondria Network Analysis tool (MiNA) run on ImageJ 1.54p as described at ^2^. Briefly, images were processed by the use of unsharp mask, enhance local contrast CLAHE, and median filtering. Then, images were binarized (on the black background), skeletonized and analyzed using MiNA Analyze Morphology plugin. More than 50 cardiomyocytes from WT and MVI-KO hearts were analyzed to describe mitochondria footprint, branch length mean and network branches mean.

### 2.6 Western blot analysis

Frozen heart ventricles were homogenized as described earlier ^19^. For assessment of OXPHOS heart ventricles were homogenized in phosphate buffer (93 mM K_2_HPO_4_, 6.5 mM KH_2_PO_4_), pH 7.2, supplemented with 1 mM PMSF at 1:50 weight-to-volume (w/v) ratio. Tissue extracts were resuspended with 5x Laemmli sample buffer, incubated for 20 min at 37°C, and stored at −25°C.

Equal amounts of samples (∼25 µg of protein/well and ∼35 µg of protein/well for OXPHOS analysis) were separated by 10-15% SDS-polyacrylamide gels (SDS-PAGE), and then transferred to a nitrocellulose membrane (Amersham). Transfer efficiency was examined by Ponceau staining of the membrane immediately after the transfer. The membranes were blocked with 3% (w/v) fat-free milk or 5% (w/v) BSA in Tris-buffered saline (TBS) containing 0.2% Triton X-100 (TBS-T) for 1 h, and then probed with the appropriate primary (Supplementary material online, *Table S4*) and secondary antibodies (Supplementary material online, *Table S5*). The level of GAPDH was used as an internal protein loading control.

The protein bands were visualized using the enhanced chemiluminescence reagent (ECL) according to the manufacturer’s instructions (Millipore, P90720). Band densitometry quantification was performed using free image processing and analysis software ImageJ 1.54p.

### 2.7 Purification of GST-fusion protein

Recombinant human myosin VI (C-terminal globular tail, GST-MVI-GD) fused to glutathione S-transferase (GST) was produced in *Escherichia coli* Rosetta (DE3)pLysS cells (Novagen) carrying the appropriate expression plasmid. Bacterial cultures were grown in LB medium until the optical density at 600 nm reached approximately 0.6. Then, protein expression was induced by the addition of 0.5 mM isopropyl-D-1-thiogalactopyranoside (Sigma-Aldrich, 11411446001), followed by incubation for 4 h at 25°C. Next, bacterial cells were collected by centrifugation at 4300 x g for 25 min. After that, bacterial pellets were resuspended in a lysis buffer containing 50 mM Tris-HCl pH 7.5, 150 mM NaCl, 1 mM EDTA, 0.1 mM PMSF, protease and phosphatase inhibitors (Roshe, 04693116 and 04906837001, respectively). Cell lysis was performed by sonication on ice using a probe sonicator (Branson Ultrasonic S.A., S-25D) (3 cycles of 10 s pulses with 20 s cooling intervals at 30% amplitude) to prevent overheating of the sample. The cell lysate was further centrifuged at 40000 × g 30 min at 4°C. The recombinant GST-tagged protein was purified using glutathione affinity chromatography on Glutathione Sepharose 4B™ resin according to the manufacturer’s protocol (Merck, GE17-0756-05).

### 2.8 GST pull-down assay

Heart from 3-month-old mice was homogenized in 10 volumes (w/v) of ice-cold lysis buffer containing 50 mM Tris-HCl (pH 7.5), 150 mM NaCl, 5% glycerol, 0.5% Triton X-100, 5 mM EDTA, 50 mM NaF, 1 mM Na₃VO₄ and 0.5 mM PMSF, supplemented with a complete protease and phosphatase inhibitor cocktails. The homogenate was clarified by centrifugation for 5 min at 500 × g, and the resulting lysate was used for pull-down analysis. To eliminate proteins that bind non-specifically to glutathione resin or GST, lysates were pre-cleared by incubation with GST-bound Glutathione Sepharose 4B beads for 2 h at 4°C with gentle rotation. The beads were subsequently removed by centrifugation at 18000 × g for 10 min at 4°C.

For the pull-down assay, approximately 20 μg of purified recombinant GST-MVI-GD construct, GST, and beads alone were immobilized on 50 μl of Glutathione Sepharose 4B beads. The resin-bound proteins were then incubated with equal amounts of the pre-cleared mouse heart lysate for 4 h at 4°C. After incubation, beads were washed extensively with the lysis buffer to remove unbound proteins. Bound protein complexes were eluted by boiling the beads in SDS sample buffer and subsequently analyzed by SDS-PAGE and immunoblotting.

### 2.9 Sample preparation and protein identification by LC-MS/MS

Samples obtained from the pull-down assay were separated by SDS–PAGE using 10% gels. Following electrophoresis, proteins were visualized by staining with Coomassie Brilliant Blue R-250. After staining, corresponding gel fragments (see Supplementary material online, *Figure S2*) were excised from the experimental lane containing the GST-MVI-GD fusion protein and the control lanes containing GST and beads alone for further LC-MS/MS analysis as described earlier ^11^.

### 2.10 Analysis of mass spectrometry data

Proteins were considered as potential myosin VI binding partners only when they were identified by at least two unique peptides with Mascot scores ≥ 50, corresponding to a statistically significant match (p ≤0.05). Furthermore, these proteins had to be detected exclusively in the experimental GST-MVI-GD sample (gel lane *iii*) and be absent from both controls: the GST sample (gel lane *ii*) and the Glutathione Sepharose 4B sample (gel lane *i*). Based on these stringent selection criteria, we identified 315 putative MVI interaction partners.

### 2.11 Statistical analysis

Results were expressed as means ±SD (standard deviation) or ± SEM (standard error of the mean) for the analysis of mitochondria fractions. The number of animals was provided in the figure legends. Statistical analyses were performed using normality and longnormality tests, and parametric Welch’s t test (for normal distribution) or non-parametric Mann-Whitney t test. Statistical significance was defined as P value provided on the graphs or legends.

### 2.12 Illustration preparing

Illustrations were created with open-source smart.servier.com (image(s) provided by Servier Medical Art (https://smart.servier.com), licensed under CC BY 4.0 (https://creativecommons.org/licenses/by/4.0/) and freepik.com (figures designed by Freepik stock content).

## 3. Results

In order to reveal involvement of MVI in mitochondria organization and function, we have employed several techniques including immunofluorescence staining, transmission electron microscopy (TEM), immunodetection as well as an affinity chromatography (pull-down assay) followed by mass spectrometry analysis. The examination was performed on heart samples derived from both WT and MVI-KO male mice at different age: newborn (P0) as well as 3 months (3m) and 12 months (12m), and on the crude cardiomyocyte culture isolated from 6-days-old (P6) mouse hearts (see *Figure 1B*).

### 3.1 MVI loss affects mitochondria distribution and morphology in the heart

Immunostaining of P0, 3m and 12m transverse heart ventricle sections for mitochondrial outer translocase protein, TOMM20, revealed that mitochondria were distributed in arrays along cardiomyocytes both in WT and MVI-KO samples (*Figure 1C).* A quantification of overall mitochondrial volume based on the TOMM20 immunofluorescence did not reveal a statistical difference between WT and MVI-KO samples (*Figure 1D*). However, a profound increase in the mitochondria presence within perinuclear area was observed for 12m MVI-KO cardiomyocytes (*Figure 1C,* see *white arrows*).

Examination of the level of TOMM20 as well mitochondrial-resident chaperon, Mortalin, did not show any changes between WT and MVI-KO samples *(Figure 1E,* and Supplementary material online, *Figure S3A*). To reveal possible mechanism(s) engaged in the above described aberrant mitochondria distribution under the MVI loss, we examined the levels of Myo19 (actin-binding motor protein) and MIRO1 (mitochondrial outer membrane Rho GTPase 1), the proteins involved in mitochondria transport, endoplasmic reticulum-mitochondria contact sites, and mitochondria cristae structure ^20–22^. A densitometric analysis did not show any statistical difference in the level of both proteins at all examined time points suggesting that neither Myo19 nor MIRO1 are associated with the changes evoked by the MVI loss (Supplementary material online, *Figure S3B*).

To get more insight into mitochondria network organization under the MVI loss, we performed analysis of the primary cardiomyocyte crude culture isolated from 6-day-old (P6) hearts. Immunostaining for TOMM20 revealed that loss of MVI leads to a change of mitochondria organization pattern with prevalence of chainmail-like morphology in MVI-KO cells (*Figure 1F,* see *red arrows*). While in WT samples this pattern constitutes for about 33% of the overall mitochondria morphology, in MVI-KO ones it raises for up to 60% (and *Figure 1G*, left graph). Also, under the MVI loss the perinuclear mitochondria clustering was more pronounced with the increase up to 17% more cells with respect to WT cells (*Figure 1F, see white arrows,* and *Figure 1G*, right graph). Additionally, by means of the Mitochondria Network Analysis tool (MiNA), we estimated the mitochondria footprint, the parameter reflecting the area covered by mitochondria within the cell. As shown in *Figure 1H*, loss of MVI caused a statistically significant decrease of the footprint value further suggesting a mitochondrial network impairment in MVI-KO cardiomyocytes.

### 3.2 MVI loss causes increase in the mitochondria mass in myocardium of 12-month-old mice

Next, TEM examination was performed to further characterize mitochondria morphology and distribution under the MVI loss. The analysis of longitudinally oriented cardiomyocytes in transverse sections of left ventricles of hearts isolated from P0, 3m and 12m mice revealed that mitochondria at all examined time points were mainly distributed in arrays between the myofibrils regardless the MVI presence (*Figure 2A*); this distribution pattern was more evident in the hearts of adult mice (see Supplementary material online, *Figure S4*).

**Figure 2.**
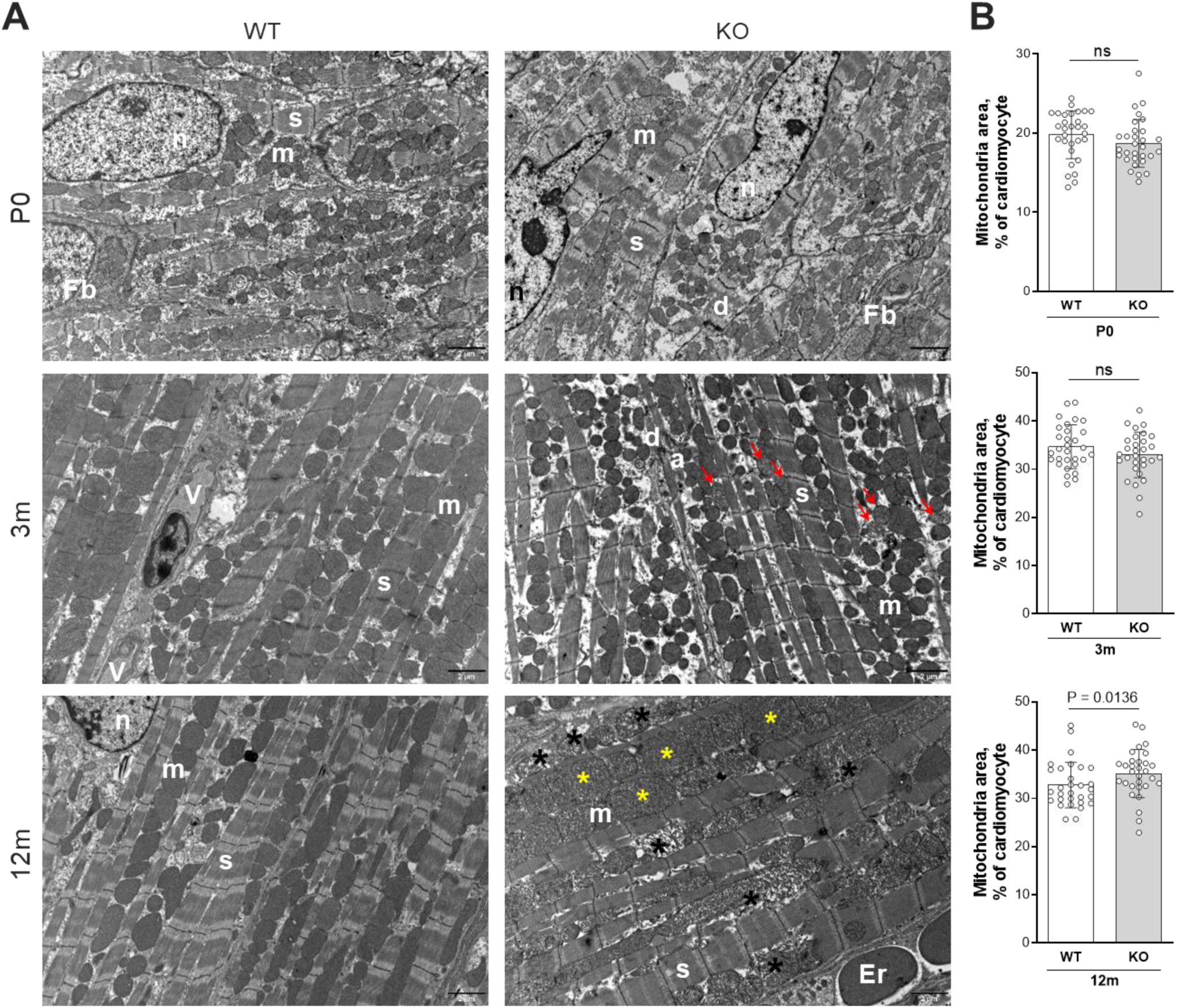
Loss of MVI leads to an age-dependent increase in mitochondria mass. **A**, representative TEM micrographs of a left ventricle in control (WT) and MVI-KO (KO) mice. Red *arrows* point to mitochondria with reduced cristae density in 3m MVI-KO mice. *Yellow asterisks* indicate large mitochondria, *black asterisks* indicate damaged mitochondria in 12m MVI-KO mice. Er, erythrocyte; Fb, fibroblast; V, vessels; n, nucleus; m, mitochondria; s, sarcomere; a, adherens junctions; d, desmosomes. Scale bar 2 µm. **B**, lack of MVI leads to a statistically significant increase in total mitochondria area in the cardiomyocytes of 12m mice. Graphs represent means ±SD. Each circle (°) represents a measurement from a single image (30 images per individual, with 3 individuals per experimental group). Statistical significance was analyzed with Welch’s t test and Mann-Whitney test, related to WT. P value is provided on the graph, ns – not statistically significant difference.

A quantification of the area covered by mitochondria (overall mitochondria area) revealed a statistically significant increase in the mitochondria mass according to the animal maturation as the mitochondrial area covered ∼20% of cardiomyocyte area in newborns and over 33% in 3m and 12m samples (*Figure 2* and *Figure S4*). While the analysis did not reveal significant changes between WT and MVI-KO samples of P0 and 3m mice, the overall mitochondria area was significantly higher in samples from 12m MVI-KO mice with respect to WT counterparts (*Figure 2B*). Also, mitochondria clustering in the perinuclear space was observed in 12m hearts confirming the results obtained with TOMM20 staining (*Figure 1C*) (Supplementary material online, *Figure S5*, see *red asterisks*). Thus, these results indicate that loss of MVI is associated with higher mitochondria accumulation and redistribution with age.

To get more insight into involvement of MVI in the maintenance of mitochondria morphology, we performed in-depth examination of TEM micrographs (see *Figure 3A-E*). We divided mitochondria in five subtypes: (i) mitochondria characteristic for cardiomyocytes (normal); (ii) mitochondria with a reduced cristae density (RCD); (iii) mitochondria with intra-cristae junctions (ICJ); (iv) donut-shaped mitochondria (DN); and (v) damaged mitochondria (DMT) (*Figure 3A*). The subtype quantification demonstrated changes in the content of particular fractions in the examined samples, especially those derived from adult mice (*Figure 3B* and Supplementary material online, *Figure S6 and S7)*. Statistically significant changes were observed in the content of RCD and DMT fractions with the increase in RCD mitochondria (about four times more than in WT) in 3m MVI-KO hearts and in DMT mitochondria (about 10 times more than in WT) in 12m MVI-KO hearts with respect to the WT counterparts. A small fraction of DN mitochondria (about 1-2%) was present only in P0 mice regardless the MVI presence (*Figure 3B*). This could be associated with a short-term exposure to hypoxia during the birth ^23^.

**Figure 3.**
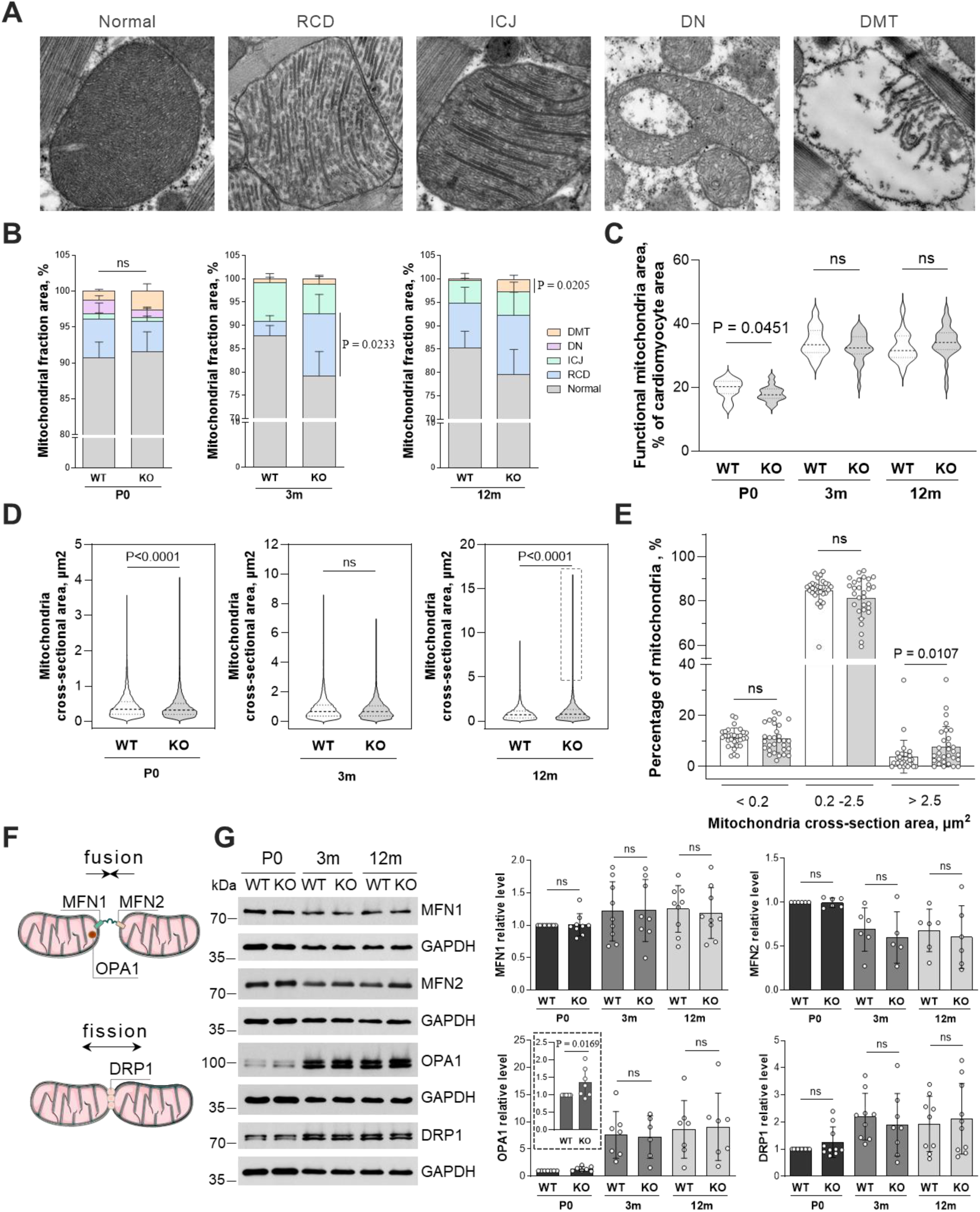
Lack of MVI leads to age-dependent changes in mitochondria morphology and dynamics. **A**, representative TEM images depicting mitochondria morphology categories analyzed in the study: normal mitochondria (normal), mitochondria with reduced cristae density (RCD), with intra-cristae junctions (ICJ), donut-shaped (DN), and damaged mitochondria (DMT). **B**, quantification of the area of individual mitochondrial fractions relative to the cardiomyocyte area (excluding cell nuclei). Results were expressed as means ± SEM. **C**, the amount of functional mitochondria (excluded DMT fraction) in cardiomyocytes. Results shown as violin plots, bold horizontal dashed line is median. **D**, evaluation of the area of individual mitochondria without fraction of DMT. Mitochondria located at the image border were not counted. Results shown as violin plots, bold horizontal dashed line representing median. *Dashed square* indicates a population of enlarged mitochondria. **E**, additionally, the cross-sectional area of individual mitochondria area from 12m mice were subdivided into 3 groups: < 0.2 µm^2^, 0.2 – 2.5 µm^2^, and > 2.5 µm^2^. Individual image analysis was indicated in the graph as (°). The results are expressed as means ±SD. Measurements at **B**, **C**, **D**, and **E** were carried out on 10 images from each of 3 different individuals (30 images in total). Statistical significance was analyzed with parametric or non-parametric t test, related to WT animals of the appropriate age. P values are provided on the graphs. ns – not statistically significant difference. **F**, schematic illustration of the mechanisms of mitochondria fusion and fission, with key proteins involved in these processes. **G**, Western Blot and densitometry of proteins involved in mitochondria dynamics. GAPDH was used as an internal loading control. The results of densitometry are expressed as means ±SD. Statistical significance was analyzed with parametric or non-parametric t test, related to WT animals of the appropriate age (P0 WT served as 1). P value is provided on the graph. ns – not statistically significant difference. Each circle on the graph (°) represents an independent biological repeat (N ≥5). *Dashed square* in the central bottom panel in **G** indicates the OPA1 level in P0 hearts on the graph with an adjusted y axis.

Analysis of the area covered by functional mitochondria (i.e. DMT excluded) revealed a statistically significant decrease in P0 MVI-KO hearts (*Figure 3C*). Interestingly, the presence of a pool of large mitochondria was observed in 12m MVI-KO heart samples (*Figure 3D*; see *dashed squares*, and Supplementary material online, *Figure S7* and *S8A*; see *dashed squares*). Therefore, we performed a detailed quantification of the mitochondria cross-section areas within 12m heart samples. The analysis demonstrated that MVI-KO hearts contained a statistically higher number of very large mitochondria (i.e. with cross-section area higher than 2.5 µm^2^) with respect to WT counterparts (*Figure 3E*). Analysis of mitochondria circularity index shoved a minimal but statistically significant decrease of a mitochondria roundness further confirming aberrations of mitochondria morphology and organization under the MVI loss (Supplementary material online, *Figure S8B*).

### 3.3 The influence of the MVI loss on the mitochondria fusion/fission in myocardium

Dynamics of mitochondria is regulated mainly by their movement and fusion/fission events, which are known to be more intense in neonatal tissues, including cardiomyocytes ^3,6^. To assess whether the above described effects of the MVI loss on mitochondria morphology in the heart are associated with impairment of mitochondria dynamics, we examined the level of proteins that are crucial for fusion (i.e. mitofusins 1 and 2, MFN1 and MFN2, and optic atrophy type 1, OPA1) and fission (dynamin-related protein 1, DRP1) (see *Figure 3F*). The Western blotting for the aforementioned proteins did not reveal statistical changes in their level in the examined samples with the exception of OPA1, level of which was statistically higher in P0 MVI-KO hearts with respect to WT counterparts (*Figure 3G)*.

### 3.4 Loss of MVI affects autophagy/mitophagy in cardiac tissue

It is known that alterations in the mitochondria quality control system, leading to aberrations in mitochondria network, are correlated with cardiac disorders ^1,6^. Mitochondrial autophagy (mitophagy) plays a crucial cardioprotective role and MVI has been shown to be involved in this multistep process (*Figure 4A*) ^8^. Therefore, we examined the effects of the MVI loss on mitophagy steps by assessment of the level of the key mitophagy marker proteins (*Figure 4B*): PINK1, Parkin, p62 and Optineurin, which label dysfunctional and damaged mitochondria as well as LC3 B form and Beclin 1, which by decoration of autophagosomes target them to lysosomes for degradation (*Figure 4A*) ^7,24,25^. The analysis showed a statistically significant decrease of Parkin, p62 and LC3 B form only in P0 MVI-KO hearts, and increase in the level of LC3 B in 3m MVI-KO hearts. However, we did not notice any change in the level of PINK, Optineurin and Beclin 1 in all the examined samples (*Figure 4B*).

**Figure 4.**
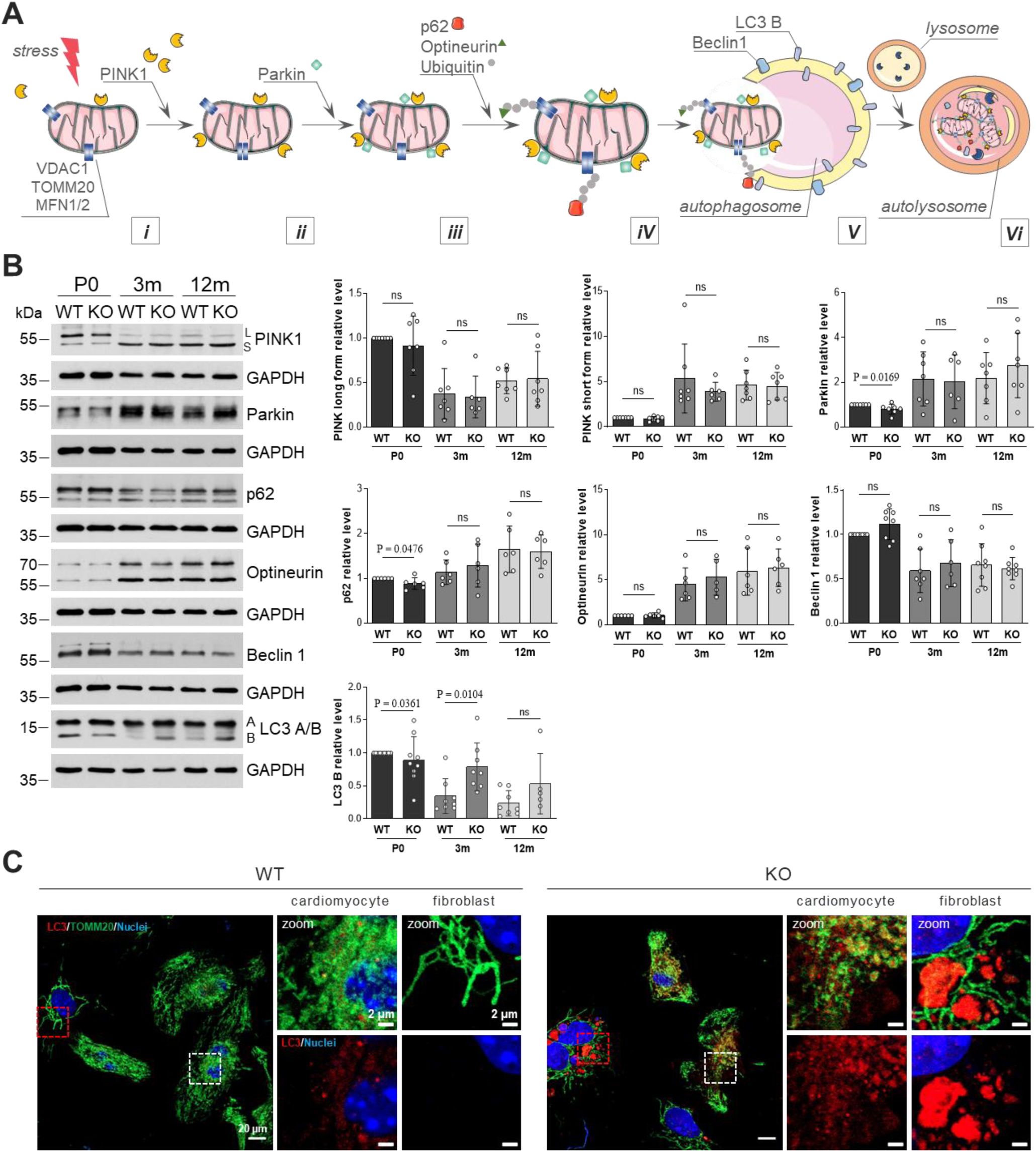
Loss of MVI impairs autophagy/mitophagy in the myocardium. **A**, Schematic diagram of mitophagy pathway. Dysfunctional and damaged mitochondria (i) are labeled by specific proteins (ii, iii, iv) and then transported to an autophagosome (v) for subsequent degradation in a lysosome (vi). **B**, representative Western blots and densitometry of the key autophagy/mitophagy-related proteins. GAPDH was used as an internal loading control. Graphs represent means ±SD. Statistical significance was analyzed with parametric or non-parametric t test, related to WT animals of the appropriate age (P0 WT served as 1). P values are provided on the graphs. ns – not statistically significant difference. Each circle (°) represents a biological repeat (N ≥ 5). **C**, double staining for autophagosomes marker LC3 and mitochondria marker TOMM20 of primary cultured cardiomyocytes and cardiac fibroblasts isolated from 6-days-old (P6) mice. *White dashed square* shows magnified cardiomyocyte fragments, *red dashed square* shows magnified fibroblast fragments.

Involvement of MVI in mitophagy was also examined by immunofluorescence staining for total LC3 in the primary crude culture of cardiomyocytes containing also cardiac fibroblasts. The analysis revealed a dramatic accumulation of large autophagosomes (marked in red) in cardiac fibroblasts and a mild increase in their presence in cardiomyocytes (*Figure 4C*). Interestingly, cardiomyocyte autophagosomes were much smaller and more dispersed within the cell in comparison with fibroblast counterparts.

### 3.5 Loss of MVI does not affect energy balance in heart ventricles

We have previously shown that loss of MVI affects mitochondria function and thus metabolism in skeletal muscles and myogenic cells ^12^. Therefore, we assayed the levels of the respiratory chain (OXPHOS) components and activity of AMPK, the energy sensor, in myocardium. Western blot analysis revealed no statistical difference in any analyzed protein between WT and MVI-KO samples in all examined time points, indicating that in the heart loss of MVI does not have such evident impact on the respiratory chain content and AMPK activity as in skeletal muscle (*Figure 5A*). However, a statistically significant decrease in the level of the active form of ribosomal protein, pS6, in P0 MVI-KO indicates that loss of MVI may impair a global protein synthesis (*Figure 5A*). Of note, S6 has been shown to interact with MVI in skeletal muscle and myogenic cells ^11,12^.

**Figure 5.**
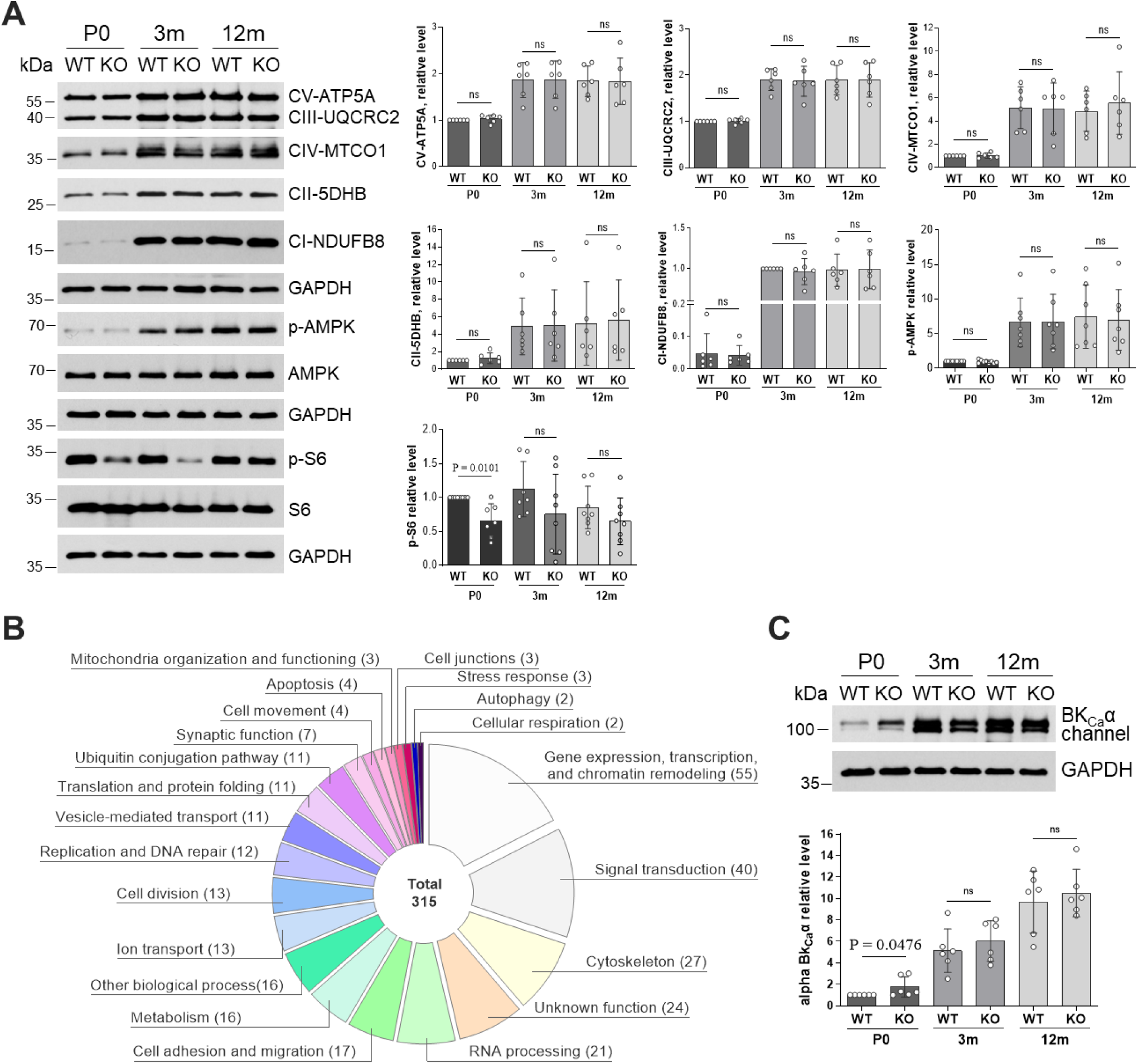
Role of MVI in key metabolic pathways. **A**, Western blot analysis and densitometry of the level of OXPHOS respiratory complex, as well as total and phosphorylated forms of AMPK and S6 (p-AMPK and p-S6). Graphs represent means ±SD. GAPDH was used as an internal loading control. Statistical significance was analyzed with parametric or nonparametric t test, related to WT animals of the appropriate age (P0 or 3m WT served as 1). P value is provided on the graph. ns – not statistically significant difference. **B**, identified potential MVI-binding protein partners in 3-month-old mouse heart. Proteins identified in MS/MS analysis were grouped according to their functions. Diagram represents the number of proteins in each group. **C**, Western blot analysis and densitometry of a potential MVI-binding partner of the BK_Ca_ channel. GAPDH was used as an internal loading control. Graph represents means ±SD. Statistical significance was analyzed with Mann-Whitney test, related to WT animals of the appropriate age (P0 WT served as 1). P value is provided on the graph. ns – not statistically significant difference. In **A** and **C** each circle (°) represents an independent experiment (N ≥ 6).

### 3.6 Identification of MVI-binding partners in mouse cardiac tissue

To further explore the effect of the MVI loss on the heart, we decided to look for potential MVI-binding partners in this organ. The analysis was performed analogously to our previous studies on other cells and tissues ^11,26,27^ with the use the pull-down assay with MVI cargo-binding domain (see *Figure 1A*) fused with glutathione-S-transferase GST (GST-MVI-GD) as a bait, followed by the glutathione-based affinity chromatography and tandem mass spectrometry (MS/MS). Homogenates of 3m WT heart ventricles were subjected to the analysis.

Three hundred and fifteen proteins were identified, which mainly based on the UniProt database (Release 2026_02) and National Library of Medicine (National Center for Biotechnology Information), have been classified into 22 functional groups (see *Figure 5B* and Supplementary material online, *Tables S6-27*). Among them were proteins already found to interact with MVI in other cells/tissues such as actin, CLIPs, DDX3X, DOCK7, histones, hnRNPs, LRPs, NUPs, TRIMs, and vinculin, making our analysis more confident ^11,27–29^. The highest number of identifications were proteins involved in gene expression, transcription, and chromatin remodeling (55), signal transduction (40) and cytoskeleton organization (27). There were also proteins engaged in RNA processing (21), cell adhesion and migration (17), metabolism (16), ion transport (13), cell division (13), replication and DNA repair (12), vesicles mediated transport (11), translation and protein folding (11), ubiquitin conjugation pathway (11), synaptic function (7), cell movement (4), apoptosis (4), mitochondria organization and functioning (3), cell junctions (3), stress response (3), autophagy (2), and cell respiration (2). There were also proteins classified into unknown functions (24) and other biological processes (16).

Similar analyses performed for other cells/tissues also revealed a number of identifications involved in a panoply of cellular functions ^11,14,26,27,29,30^. What is different from those studies is identification in heart homogenates a number of proteins involved in the processes related to mitochondria organization and function, cellular respiration, and ion transport (Supplementary material online, *Table S6, S8, S11*). Among them were a constituent of respiratory channel complex II (succinate dehydrogenase assembly factor 2, *Sdhaf1*), and alpha subunit of calcium-activated large conductance potassium (BK_Ca_) channel (*Kcnma1*) that can be also targeted to the inner mitochondrial membrane ^31,32^. While we showed herein that the level of complex II was not affected by the MVI loss (*Figure 5A*), there is no information on dependence of the level of the BK_Ca_ channel on the presence or absence of MVI. Therefore, we performed Western blot analysis for the levels of BK_Ca_ channel in P0, 3m and 12m WT and MVI-KO hearts. The level of BK_Ca_ alpha subunit protein was significantly increased only in P0 MVI-KO hearts with respect to WT counterparts (*Figure 5C*), suggesting a potential functionality of MVI-BK_Ca_ channel *in vivo* during heart development and maturation.

## 4. Discussion

The study presented herein points at unconventional MVI as an important regulator of the mitochondria maintenance. Our results obtained from the hearts of newborn (P0) and adult (3m and 12m) mice, as well as on primary cardiomyocyte culture, indicate that MVI is involved in processes associated with mitochondria quality control and network organization. We found that this involvement varies with age, as the effects observed in the hearts of newborn MVI-KO mice result in increased mitochondria mass and amounts of impaired mitochondria in 12m hearts. We believe that this accumulation of non-functional mitochondria, as well as alterations in the structure and distribution of mitochondria, contribute to the age-progressing cardiac remodeling and dysfunction that we have recently reported ^19^. Mitochondria in both WT and MVI-KO hearts at all examined lifetime points are arranged in longitudinal arrays between the myofibrils. A more pronounced perinuclear distribution was observed, specifically, in 12m MVI-KO hearts. The profound effect of the MVI loss on mitochondrial network organization was also observed in isolated cardiomyocytes, where a chainmail-like network was predominant, further confirming an involvement of MVI in the mitochondria maintenance. This mitochondria clustering/accumulation in the perinuclear space could be an effect of the adaptation of mitochondria network to metabolic demand as perinuclear mitochondria have been suggested to produce ATP and regulate local Ca^2+^ flow closely to the nucleus, thereby influencing nuclear functions ^3^. Also, it was showed that in adult cardiomyocytes, intrafibrillar mitochondria trafficking is limited by a tightly packed contractile apparatus ^3^. Interestingly, our data showed that the level of Myo19 (another unconventional myosin specifically involved in mitochondria translocation and positioning) decreases with age (though this decrease was not associated with the MVI loss) and overall mitochondria mass increases in adult mice seems to confirm this notion. It is also known that mitochondria localized to the perinuclear region could be functionally altered in failing cardiomyocytes where depressed Ca^2+^ uptake, decreased mitochondrial membrane potential and increased ROS generation have been reported ^33^. This is in line with our results showing that loss of MVI caused a statistically significant increase in the overall mitochondria mass as well as an accumulation of large (i.e. with cross-section area >2.5 µm^2^) functional mitochondria with the concomitant increase in the presence of damaged mitochondria in 12m MVI-KO hearts. It is noteworthy that while the amount of damaged mitochondria was not changed in 3m MVI-KO hearts, the amount of mitochondria with reduced cristae density was increased at this lifetime point, indicating that the process of mitochondria failing has already started.

We know from the studies on myogenic cells that either deficiency or loss of MVI causes mitochondria dysfunction, leading to lower respiratory capacity and reduced ATP production accompanied by an increase in activation (phosphorylation) of AMPK, the energy sensor, as shown in recent research by ^12^. It is noteworthy that changes in mitochondria functioning were also detected in the soleus muscle of MVI-KO adult mice and newborns ^12^. Intriguingly, the data presented herein are distinct from those gathered on the skeletal muscle and myogenic cells as we did not observe any effect of the MVI loss on the levels of active AMPK and components of the respiratory chain indicating that oxidative phosphorylation and energy balance were not significantly altered. However, similar to skeletal muscle and neurosecretory PC12 cells our data indicate that loss of MVI could affect the overall protein synthesis reflected in the reduction of the level of active (phosphorylated) S6 protein in MVI-KO P0 hearts ^12,34^. Importantly, this tendency is also observed in the hearts of adult MVI-KO mice. We believe that a decrease in the S6 phosphorylation can results from a downregulation of Akt/mTOR pathway which we observed in P0 MVI-KO hearts ^19^.

Thus, our results indicate that while some of the MVI-associated mechanisms could be general for metabolic processes, the other ones, particularly the ones associated with mitochondria function, could be distinct and specific (perhaps cardioprotective?) to the heart. One of such mechanism may be associated with the large-conductance calcium-activated potassium channel, BK_Ca_, which is located on the plasma membrane of cardiomyocytes during early stages of mouse development, and it is also present in the inner mitochondrial membrane in adults ^31,32^. This channel participates in a variety of vital processes, such as gene expression, vascular tone, neuronal excitability, and cardiac rhythmicity. Furthermore, its inhibition increases risk of myocardial infarction in neonatal murine and rat cardiomyocytes, whereas its activation provides cardioprotection in adult animals ^32^. We observed that the loss of MVI results in a statistically significant increase in the BK_Ca_ level in newborn MVI-KO hearts, indicating that MVI deficiency may be compensated by an increased channel synthesis. It is noteworthy that BK_Ca_ channel expression and function has been reported to be altered under conditions of chronic oxidative stress, hypertension, cardiac hypertrophy, and hypoxic conditions ^35^. Importantly, Snell’s waltzer mice exhibit both hypertension and cardiac hypertrophy, which occur already at the embryonic stage ^19,36,37^. These observations seem to indicate the potential involvement of BK_Ca_ in the MVI-loss-evoked changes in the heart. We have found that MVI likely interacts with BK_Ca_ by identification of this channel as a putative MVI binding partner. It is noteworthy that several proteins involved in mitochondria function, respiration and ion transport have been also found among the potential MVI partners including: mitochondrial protein AarF domain-containing protein kinase 1, ADCK1 (involved in the mitochondria cristae formation, fusion/fission dynamics, ATP and ROS production, and mitochondria homeostasis), succinate dehydrogenase assembly factor 2 (component of the respiratory complex II), L-lactate dehydrogenase, LDH (involved in the maintenance of ATP production under low-oxygen conditions), and voltage-dependent calcium channel gamma-8 subunit (involved in modulation of high-voltage activated calcium channels activity) ^38–40^. These findings suggest potential novel mechanisms of MVI involvement in the mitochondria maintenance and function in the heart. However, without further studies confirming the nature of these interactions and their functionality, this hypothesis remains so far speculative.

Besides a possible involvement of BK_Ca_ (and other putative partners) in the MVI-loss-associated changes, we believe that our results point at two major mechanisms behind the observed phenotype. One is related to mitochondria dynamics, namely fusion, and the second is connected with mitochondria quality control, namely mitophagy. Interestingly, the effects of the MVI loss on these two processes seems to be more pronounced in the hearts of newborn MVI-KO mice. From the proteins involved in mitochondria dynamics that we tested (i.e. Drp1, MFN1 and MFN2, and OPA1), only OPA1, involved in the fusion, was affected by the MVI loss as its level was increased in MVI-KO hearts and only in newborns. In our opinion, this increase reflects more intensive fusion events at the early stage of mouse life that may later contribute to an increase in mitochondria mass and formation of large mitochondria in adults. A similar effect of the MVI loss was observed in mouse embryonic fibroblasts, MEFs, derived from Snell’s waltzer mice (*Myo6^sv/sv^*) where a ∼50% increase in mitochondria mass was detected by flow cytometry analysis ^8^.

It has been shown that MVI localizes to mitochondria both in non-muscle cells and in skeletal muscle and myogenic cells ^8,12^. It have been reported that MVI plays a key role in mitophagy by its involvement in recruitment of selectively marked damaged mitochondria to autophagosomes, through its interaction with Parkin, and encapsulating the damaged mitochondria in actin-rich cages ^8^. We showed that the level of proteins involved in mitophagy, i.e. Parkin, p62 and LC3 B form was decreased in the hearts of P0 MVI-KO mice. However, the level of LC3 B form (involved in mitophagosome formation) was significantly increased in adult, 3m MVI-KO mice. These data indicate that in the MVI-KO hearts alterations in the labeling of damaged mitochondria are already occurring in newborns, and impairment of autophagosomes transport or/and utilization in lysosomes are disclosed in adult mice. Interestingly, these observations are in line with data by ^3^ who reported that intra-fibrillar mitochondria could move toward the perinuclear region upon mitochondria damage to participate in mitophagy and, as it has been mentioned above, we observed a significant accumulation of mitochondria next to the nuclei. Also, we observed a mild auto(mito)phagosome formation (via LC3 immunostaining) in our primary crude cardiomyocyte culture containing fibroblasts. The use of such culture enabled us to observe a different autophagosome formation pattern in the two cell types in the same area of view. We are the first to show that autophagosomes formed in MVI-KO cardiomyocytes are much smaller, more uniform in size, and more dispersed within the cell in comparison with fibroblast counterparts, once again confirming the distinct (and yet not revealed) mechanisms occurring in the hearts. These data are in line with the previous studies on non-muscle cells showing that loss of MVI evokes alterations in subsequent degradation of mitophagosomes ^8,41^.

## Authors’ contributions

O.K.M., A.S., and M.J.R. designed research. O.K.M., L.L., J.N., and W.F.K performed experiments. K.S. performed mass spectrometry analysis. O.K.M., L.L., J.N., and D.W. analysed experimental results. O.K.M., J.N. and M.J.R. interpreted results of experiments. O.K.M. wrote original draft of manuscript. O.K.M. prepared figures. O.K.M. and M.J.R. edited the final manuscript with input and approval from all authors. O.K.M. and M.J.R. provided supervision. All authors read and approved the manuscript.

## Acknowledgements

We thank Prof. Folma Buss from the Cambridge Institute for Medical Research, University of Cambridge, United Kingdom for her gift of Snell’s waltzer mice. Also, we thank Dr Volodymyr Balatskyi from the Laboratory of Molecular Medical Biochemistry from the Nencki Institute for sharing their expertise in the cardiomyocyte isolation and culture. TEM imaging was performed at the Laboratory of Electron Microscopy, which serves as an imaging core facility at the Nencki Institute of Experimental Biology, and is a part of the infrastructure of the Polish Euro-BioImaging Node. Polish Node is supported by the project co-financed by the Minister of Education and Science based on contract No 2022/WK/05 (Polish Euro-BioImaging Node “Advanced Light Microscopy Node Poland”). Confocal imaging was performed at the Laboratory of Imaging Tissue Structure and Function which serves as an imaging core facility at the Nencki Institute of Experimental Biology and is a part of the infrastructure of the Polish Euro-BioImaging Node.

## References

1. Ramaccini D, Montoya-Uribe V, Aan FJ, Modesti L, Potes Y, Wieckowski MR, Krga I, Glibetić M, Pinton P, Giorgi C, Matter ML. Mitochondrial Function and Dysfunction in Dilated Cardiomyopathy. Front Cell Dev Biol. 2021;8:624216.

2. Shamseldin HE, Alhashem A, Tabarki B, Abdulwahab F, Hashem M, Sougrat R, Alkuraya FS. Mitochondrial “dysmorphology” in variant classification. Hum Genet. 2022;141(1):55–64.

3. Lu X, Thai PN, Lu S, Pu J, Bers DM. Intrafibrillar and Perinuclear Mitochondrial Heterogeneity in Adult Cardiac Myocytes. J Mol Cell Cardiol. 2019;136:72–84.

4. Li X, Jiang O, Chen M, Wang S. Mitochondrial homeostasis: shaping health and disease. Curr Med. 2024;3(1):5.

5. Valente AJ, Maddalena LA, Robb EL, Moradi F, Stuart JA. A simple ImageJ macro tool for analyzing mitochondrial network morphology in mammalian cell culture. Acta Histochem. 2017;119(3):315–326.

6. Li A, Gao M, Jiang W, Qin Y, Gong G. Mitochondrial Dynamics in Adult Cardiomyocytes and Heart Diseases. Front Cell Dev Biol. 2020;8.

7. Yang H, Zhang L, Qian H. The dual regulation of mitophagy in myocardial injury: From molecular mechanisms to targeted therapies. Tissue Cell. 2026;99:103318.

8. Kruppa AJ, Kishi-Itakura C, Masters TA, Rorbach JE, Grice GL, Kendrick-Jones J, Nathan JA, Minczuk M, Buss F. Myosin VI-Dependent Actin Cages Encapsulate Parkin-Positive Damaged Mitochondria. Dev Cell. 2018;44(4):484–499.e6.

9. Wen H, Deng H, Li B, Chen J, Zhu J, Zhang X, Yoshida S, Zhou Y. Mitochondrial diseases: from molecular mechanisms to therapeutic advances. Signal Transduct Target Ther. 2025;10(1):9.

10. Pavlović N, Todorović P, Maglica M, Kumrić M, Vukojević K, Pogorelić Z, Božić J. Mitochondrial Dysfunction in Cardiomyopathy and Heart Failure: From Energetic Collapse to Therapeutic Opportunity. Biomolecules. 2025;15(11):1572.

11. Karolczak J, Sobczak M, Skowronek K, Rędowicz MJ. A Kinase Anchoring Protein 9 Is a Novel Myosin VI Binding Partner That Links Myosin VI with the PKA Pathway in Myogenic Cells. BioMed Res Int. 2015;2015(1):816019.

12. Wojton D, Dymkowska D, Matysniak D, Topolewska M, Redowicz MJ, Lehka L. Unconventional myosin VI is involved in regulation of muscle energy metabolism. Am J Physiol-Cell Physiol. 2025;329(4):C1004–C1021.

13. Sweeney HL, Houdusse A. Myosin VI Rewrites the Rules for Myosin Motors. Cell. 2010;141(4):573–582.

14. Magistrati E, Polo S. Myomics: myosin VI structural and functional plasticity. Curr Opin Struct Biol. 2021;67:33–40.

15. Deol MS, Green MC. Snell’s waltzer, a new mutation affecting behaviour and the inner ear in the mouse. Genet Res. 1966;8(3):339–345.

16. Di Benedetto G, Gerbino A, Lefkimmiatis K. Shaping mitochondrial dynamics: The role of cAMP signalling. Biochem Biophys Res Commun. 2018;500(1):65–74.

17. Arden SD, Tumbarello DA, Butt T, Kendrick-Jones J, Buss F. Loss of cargo binding in the human myosin VI deafness mutant (R1166X) leads to increased actin filament binding. Biochem J. 2016;473(19):3307–3319.

18. Lam J, Katti P, Biete M, Mungai M, AshShareef S, Neikirk K, Garza Lopez E, Vue Z, Christensen TA, Beasley HK, Rodman TA, Murray SA, Salisbury JL, Glancy B, Shao J, Pereira RO, Abel ED, Hinton A Jr. A Universal Approach to Analyzing Transmission Electron Microscopy with ImageJ. Cells. 2021;10(9):2177.

19. Karatsai O, Lehka L, Wojton D, Grabowska AI, Duda MK, Lenartowski R, Redowicz MJ. Unconventional myosin VI in the heart: Involvement in cardiac dysfunction progressing with age. Biochim Biophys Acta Mol Basis Dis. 2023;1869(6):166748.

20. Covill-Cooke C, Kwizera B, López-Doménech G, Thompson CO, Cheung NJ, Cerezo E, Peterka M, Kittler JT, Kornmann B. Shared structural features of Miro binding control mitochondrial homeostasis. EMBO J. 2024;43(4):595–614.

21. Quintero OA, DiVito MM, Adikes RC, Kortan MB, Case LB, Lier AJ, Panaretos NS, Slater SQ, Rengarajan M, Feliu M, Cheney RE. Human Myo19 is a novel myosin that associates with mitochondria. Curr Biol CB. 2009;19(23):2008–2013.

22. Attia A, Majstrowicz K, Shembekar S, Honnert U, Nikolaus P, Lohmann B, Bähler M. Myosin-19 and Miro Regulate Mitochondria-Endoplasmic Reticulum Contacts and Mitochondria Inner Membrane Architecture. Cells. 2025;14(21):1657.

23. Long Q, Zhao D, Fan W, Yang L, Zhou Y, Qi J, Wang X, Liu X. Modeling of Mitochondrial Donut Formation. Biophys J. 2015;109(5):892–899.

24. Zhu CL, Yao RQ, Li LX, Li P, Xie J, Wang JF, Deng XM. Mechanism of Mitophagy and Its Role in Sepsis Induced Organ Dysfunction: A Review. Front Cell Dev Biol. 2021;9.

25. D’Arcy MS. Mitophagy in health and disease. Molecular mechanisms, regulatory pathways, and therapeutic implications. Apoptosis. 2024;29(9):1415–1428.

26. Karolczak J, Sobczak M, Majewski L, Yeghiazaryan M, Jakubiec-Puka A, Ehler E, Sławińska U, Wilczyński GM, Rędowicz MJ. Myosin VI in skeletal muscle: its localization in the sarcoplasmic reticulum, neuromuscular junction and muscle nuclei. Histochem Cell Biol. 2013;139(6):873–885.

27. Majewski L, Nowak J, Sobczak M, Karatsai O, Havrylov S, Lenartowski R, Suszek M, Lenartowska M, Redowicz MJ. Myosin VI in the nucleus of neurosecretory PC12 cells: Stimulation-dependent nuclear translocation and interaction with nuclear proteins. Nucleus. 2018;9(1):125–141.

28. Wollscheid HP, Biancospino M, He F, Magistrati E, Molteni E, Lupia M, Soffientini P, Rottner K, Cavallaro U, Pozzoli U, Mapelli M, Walters KJ, Polo S. Diverse functions of myosin VI elucidated by an isoform-specific α-helix domain. Nat Struct Mol Biol. 2016;23(4):300–308.

29. Finan D, Hartman MA, Spudich JA. Proteomics approach to study the functions of Drosophila myosin VI through identification of multiple cargo-binding proteins. Proc Natl Acad Sci U S A. 2011;108(14):5566–5571.

30. Tumbarello DA, Kendrick-Jones J, Buss F. Myosin VI and its cargo adaptors - linking endocytosis and autophagy. J Cell Sci. 2013;126(Pt 12):2561–2570.

31. Szabo I, Szewczyk A. Mitochondrial Ion Channels. Annu Rev Biophys. 2023; 52:229–254.

32. Sanghvi S, Szteyn K, Ponnalagu D, Sridharan D, Lam A, Hansra I, Chaudhury A, Majumdar U, Kohut AR, Gururaja Rao S, Khan M, Garg V, Singh H. Inhibition of BK_Ca_ channels protects neonatal hearts against myocardial ischemia and reperfusion injury. Cell Death Discov. 2022;8(1):175.

33. Voglhuber J, Holzer M, Radulović S, Thai PN, Djalinac N, Matzer I, Wallner M, Bugger H, Zirlik A, Leitinger G, Dedkova EN, Bers DM, Ljubojevic-Holzer S. Functional remodelling of perinuclear mitochondria alters nucleoplasmic Ca^2+^ signalling in heart failure. Philos Trans R Soc B Biol Sci. 2022;377(1864):20210320.

34. Nowak J, Lenartowski R, Kalita K, Lehka L, Karatsai O, Lenartowska M, Rędowicz MJ. Myosin VI in the nucleolus of neurosecretory PC12 cells: its involvement in the maintenance of nucleolar structure and ribosome organization. Front Physiol. 2024;15:1368416.

35. Ochoa SV, Otero L, Aristizabal-Pachon AF, Hinostroza F, Carvacho I, Torres YP. Hypoxic Regulation of the Large-Conductance, Calcium and Voltage-Activated Potassium Channel, BK. Front Physiol. 2021;12:780206.

36. Gotoh N, Yan Q, Du Z, Biemesderfer D, Kashgarian M, Mooseker MS, Wang T. Altered renal proximal tubular endocytosis and histology in mice lacking myosin-VI. Cytoskelet Hoboken NJ. 2010;67(3):178–192.

37. Hegan PS, Lanahan AA, Simons M, Mooseker MS. Myosin VI and cardiomyopathy: Left ventricular hypertrophy, fibrosis, and both cardiac and pulmonary vascular endothelial cell defects in the Snell’s waltzer mouse. Cytoskelet Hoboken NJ. 2015;72(8):373–387.

38. Jacquet N, Zhao Y. The ADCK Kinase Family: Key Regulators of Bioenergetics and Mitochondrial Function and Their Implications in Human Cancers. Int J Mol Sci. 2025;26(12):5783.

39. Flood D, Lee ES, Taylor CT. Intracellular energy production and distribution in hypoxia. J Biol Chem. 2023;299(9):105103.

40. Yang L, Katchman A, Morrow JP, Doshi D, Marx SO. Cardiac L-type calcium channel (Cav1.2) associates with gamma subunits. FASEB J Off Publ Fed Am Soc Exp Biol. 2011;25(3):928–936.

41. Tumbarello DA, Waxse BJ, Arden SD, Bright NA, Kendrick-Jones J, Buss F. Autophagy-receptors link myosin VI to autophagosomes to mediate Tom1-dependent autophagosome maturation and fusion with the lysosome. Nat Cell Biol. 2012;14(10):1024–1035.

