## Supplementary material for "Loss of myosin VI leads to age-dependent changes in the maintenance of heart ventricle mitochondria"

**Supplementary Table S1.** List of primers used for genotyping.

| Primer name | Primer sequence |
|---|---|
| SV1 | 5′-CTGACCCTGATCACTTAGCAGAGTTG-3′ |
| SV2 | 5′-CATTGGGCCAGGTCACAGAAGTAAGC-3′ |
| SV3 | 5′-GGTCCTCTGAAAGAGTAACC-3′ |

**Supplementary Table S2.** List of primary antibodies used in the immunocytochemical staining.

| № | Antigen | Manufacturer/ Catalog number | Dilutions | Conditions |
|---|---|---|---|---|
| 1 | α-Actinin | Abcam/ab9465 | 1:75 | Overnight at 4°C |
| 2 | LC3 A/B | Cell Signaling Technology/83506S | 1:25 | Overnight at 4°C |
| 3 | TOMM20 | Abcam/ab186735 | 1:50 | Overnight at 4°C |
| 4 | Vimentin | Abcam/ab92547 | 1:75 | Overnight at 4°C |

**Supplementary Table S3.** List of secondary antibodies used in the immunocytochemical staining.

| № | Antigen | Manufacturer/ Catalog number | Dilutions | Conditions |
|---|---|---|---|---|
| 1 | Alexa Fluor 488-conjugated anti-rabbit | Invitrogen/A11008 | 1:1000 | 1.5 h at RT |
| 2 | Alexa Fluor 546-conjugated anti-mouse | Invitrogen/A11003 | 1:1000 | 1.5 h at RT |

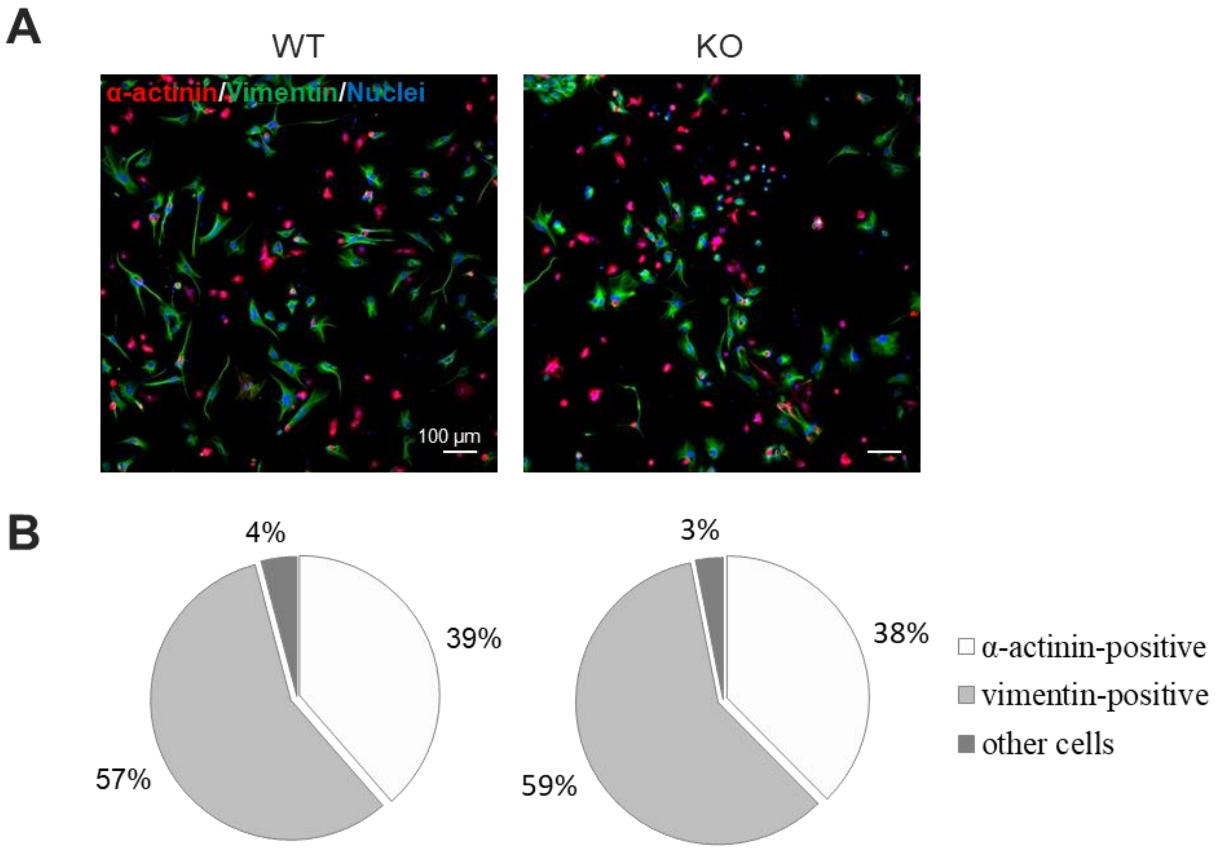


**Supplementary Figure S1.** Primary crude culture of cardiomyocytes containing cardiac fibroblasts isolated from 6-days-old mice. **A**, immunofluorescence staining for the cardiomyocyte-specific marker α-actinin and fibroblast-specific marker, vimentin. Nuclei were labeled with DAPI. **B**, analysis of cardiomyocyte culture purity. Pie chart graph showing a percentage of major cell populations within the culture.

**Supplementary Table S4.** List of primary antibodies used in Western blot analysis.

| № | Antigen | Manufacturer/ Catalog number | Dilutions/Conditions | | |
|---|---|---|---|---|---|
| 1 | AMPKα | Cell Signaling Technology/2532S | 1:1500 | 3% milk | Overnight at 4°C |
| 2 | p-AMPKα (T172) | Cell Signaling Technology/2535S | 1:1500 | 5% BSA | Overnight at 4°C |
| 3 | Beclin 1 | Cell Signaling Technology/3738S | 1:1000 | 3% milk | Overnight at 4°C |
| 4 | $BK_{Ca}$ α channel | Abcam/ab192759 | 1:1000 | 3% milk | Overnight at 4°C |
| 5 | DRP1 | Cell Signaling Technology/8570s | 1:3000 | 3% milk | Overnight at 4°C |
| 6 | GAPDH | Millipore/MAB374 | 1:15000 | 3% milk | 1 h at RT |
| 7 | GRP75/mortalin | Cell Signaling Technology/2816S | 1:200 | 3% milk | Overnight at 4°C |
| 8 | LC3 A/B | Cell Signaling Technology/4108S | 1:750 | 3% milk | Overnight at 4°C |
| 9 | MFN1 | Abcam/ab221661 | 1:2000 | 3% milk | Overnight at 4°C |
| 10 | MFN2 | Cell Signaling Technology/9482S | 1:2000 | 3% milk | Overnight at 4°C |
| 11 | MIRO1 | Antibodies/A309670 | 1:500 | 3% milk | Overnight at 4°C |
| 12 | MVI | Proteus/25-6791 | 1:1000 | 3% milk | Overnight at 4°C |
| 13 | Myo19 | Proteintech/23906-1-AP | 1:300 | 3% milk | Overnight at 4°C |
| 14 | OPA1 | Cell Signaling Technology/80471S | 1:5000 | 3% milk | Overnight at 4°C |
| 15 | Optineurin | Santa Cruz Biotechnology/sc-166576 | 1:1000 | 3% milk | Overnight at 4°C |
| 16 | OXPHOS | Abcam/ab110413 | 1:3000/ 1:200 | 3% milk | Overnight at 4°C |
| 17 | p62 | Cell Signaling Technology/5114S | 1:400 | 3% milk | Overnight at 4°C |
| 18 | Parkin | Cell Signaling Technology/2132S | 1:250 | 3% milk | Overnight at 4°C |
| 19 | PINK1 | Abclonal/A7131 | 1:500 | 3% milk | Overnight at 4°C |
| 20 | S6 | Cell Signaling Technology/2217S | 1:3000 | 3% milk | Overnight at 4°C |
| 21 | p-S6 (S235/236) | Cell Signaling Technology/4858S | 1:3000 | 5% BSA | Overnight at 4°C |
| 22 | TOMM20 | Abcam/ab186735 | 1:1000 | 3% milk | Overnight at 4°C |

**Supplementary Table S5.** List of secondary antibodies used in Western blot analysis.

| № | Antigen | Manufacturer/ Catalog number | Dilutions/Conditions | | |
|---|---|---|---|---|---|
| 1 | Goat anti-mouse IgG-HRP | Millipore/AP308P | 1:5 000 -10 000 | 3% milk or 5% BSA | 1 h at RT |
| 2 | Goat anti-rabbit IgG-HRP | Millipore/AP307P | 1:5 000 -10 000 | 3% milk or 5% BSA | 1 h at RT |

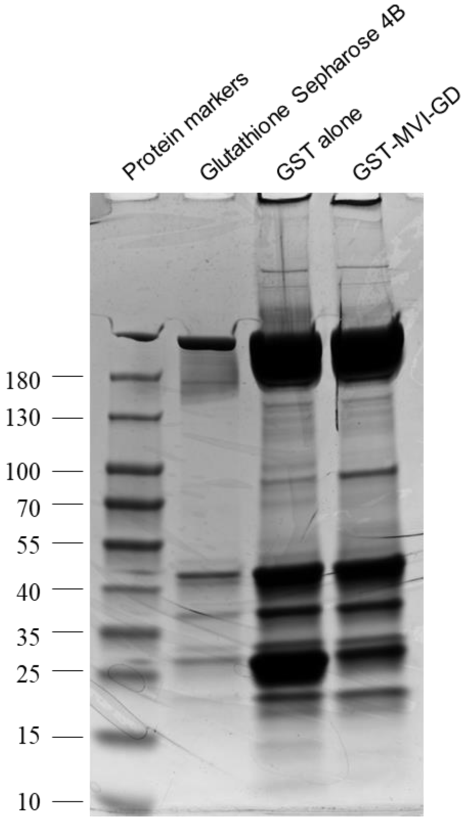


**Supplementary Figure S2.** Screening for potential MVI protein partners in the mouse heart using a pull-down assay. Homogenates obtained from the heart of 3-month-old WT mice bound to a Glutathione Sepharose 4B resin, the resin with GST alone or the resin with MVI globular tail domain fused with GST (GST-MVI-GD) as described in the "Methods" sections 2.8-2.10.

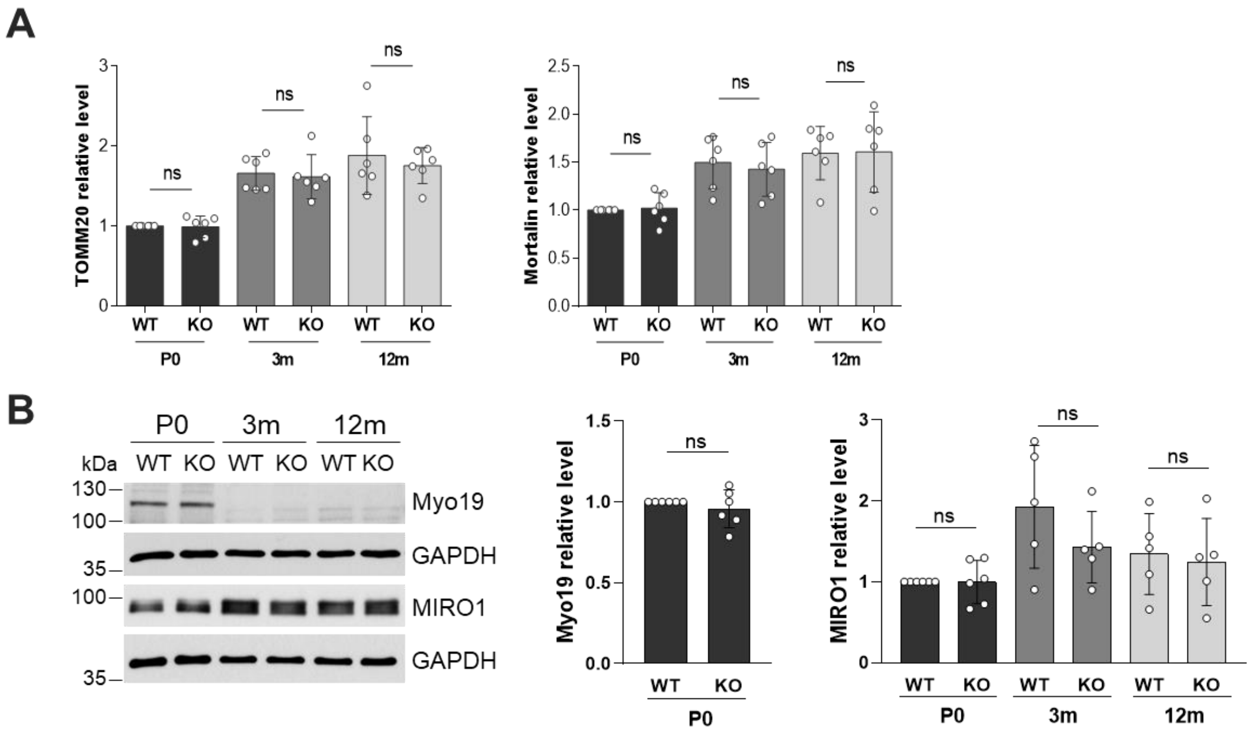


**Supplementary Figure S3.** Analysis of the levels of mitochondria-resident proteins and proteins involved in mitochondria transport. **A**, Densitometry of Western blots of mitochondria markers TOMM20 and Mortalin in newborn (P0), adult 3-month-old (3m) and 12-month-old (12m) WT and MVI-KO mice. **B**, Western blot analysis and densitometric quantification of the levels of motor protein, Myo19, and mitochondrial membrane receptor MIRO1. GAPDH was used as an internal loading control. In **A** and **B** statistical significance was analyzed with parametric or non-parametric t test, related to WT animals of the appropriate age (P0 WT served as 1). ns – not statistically significant difference. Each circle (°) represents a biological repeat (N ≥ 5).

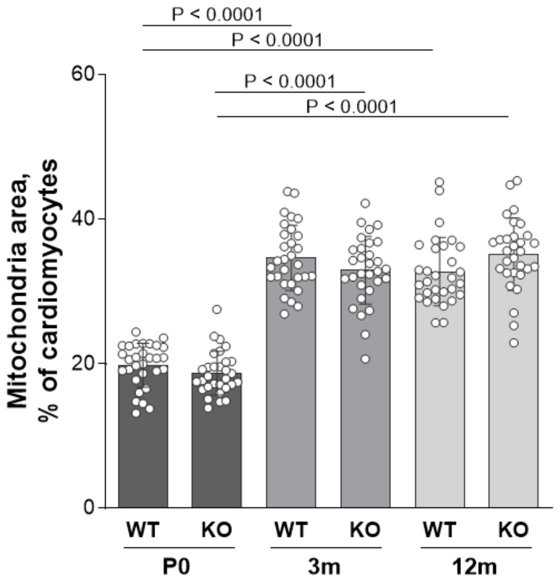


**Supplementary Figure S4.** Statistical analysis of the changes in total mitochondria area with age. Graph represent means ±SD. Each circle (°) represents a measurement from an individual TEM image (30 images from 3 different individuals in each experimental group). Statistical significance was analyzed with parametric or non-parametric t test. P values are provided on the graph.

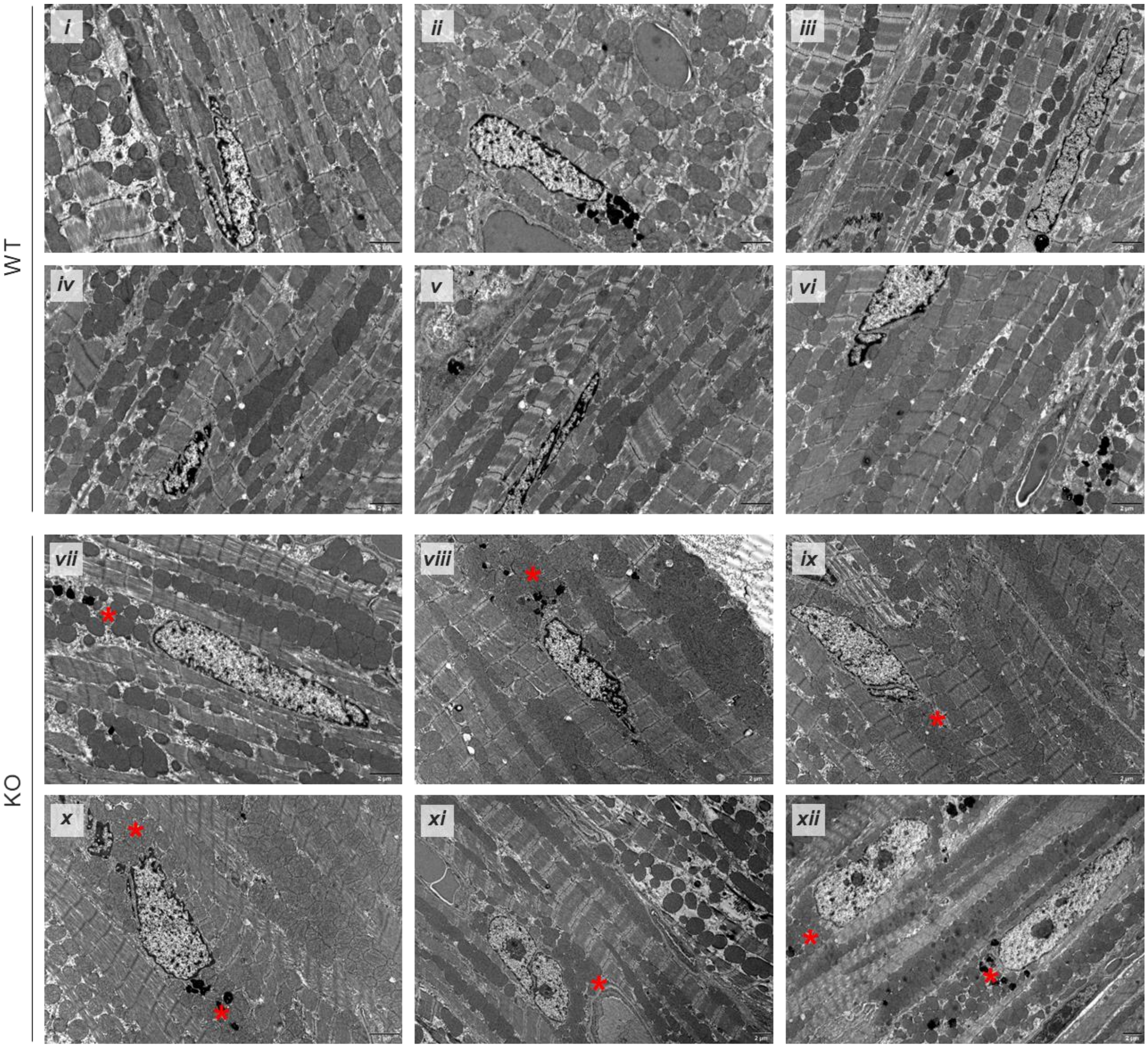


**Supplementary Figure S5.** Ultrastructural examination of perinuclear localization of the mitochondria in cardiomyocytes from 12m mice. Representative TEM micrographs of a left ventricle in control (WT; panels i-vi) and MVI-KO (KO; panels vii-xii) mice. *Red asterisks* mark perinuclear mitochondria. Scale bar, 2 μm.

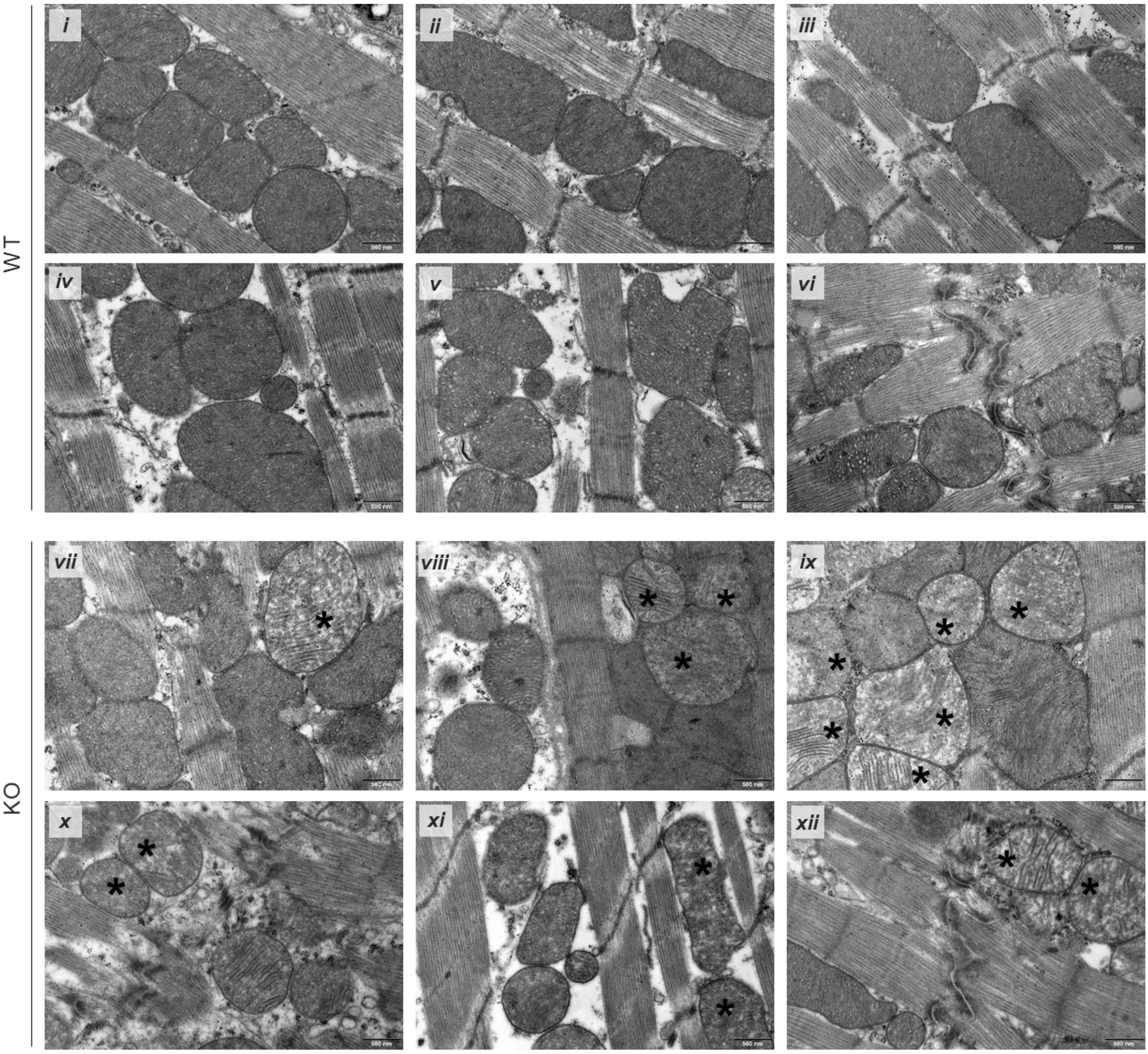


**Supplementary Figure S6.** Ultrastructural examination of mitochondria morphology in cardiomyocytes from 3m mice. Representative TEM micrographs of left ventricle in control (WT; panels i-vi) and MVI-KO (KO; panels vii-xii) mice. *Black asterisks* mark mitochondria with reduced cristae density. Scale bar, 500 nm.

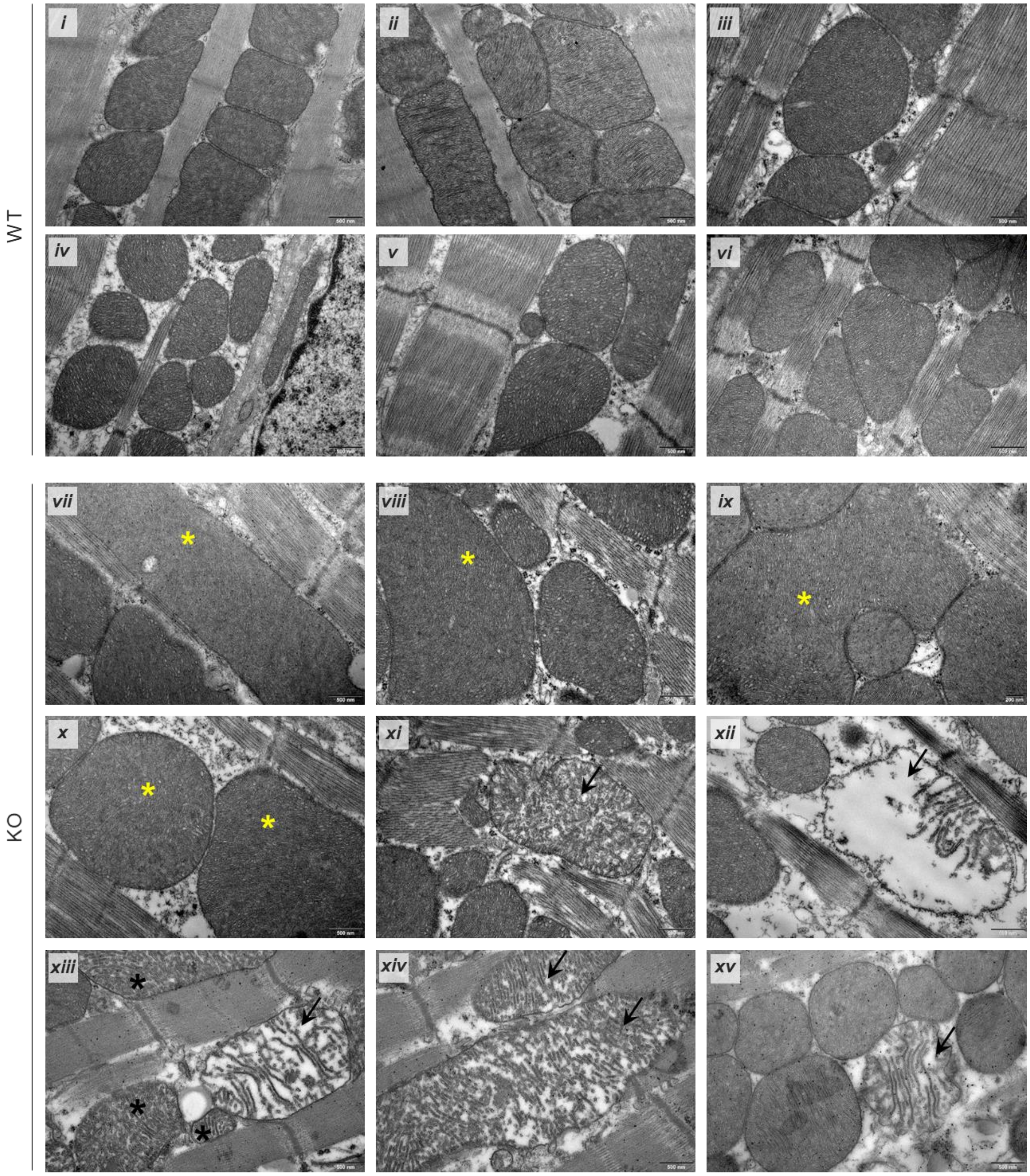


**Supplementary Figure S7.** Ultrastructural examination of mitochondria morphology in cardiomyocytes from 12m mice. Representative TEM micrographs of left ventricle in control (WT; panels i-vi) and MVI-KO (KO; panels vii-xv) mice. Micrographs in panels iii and xii were used as representative in Figure 3A for depiction of mitochondria morphology categories analyzed in the study. *Yellow asterisks* mark large mitochondria, *black asterisks* mark mitochondria with reduced cristae density, and *black asterisks* point to damaged dysfunctional mitochondria. Scale bar, 500 nm.

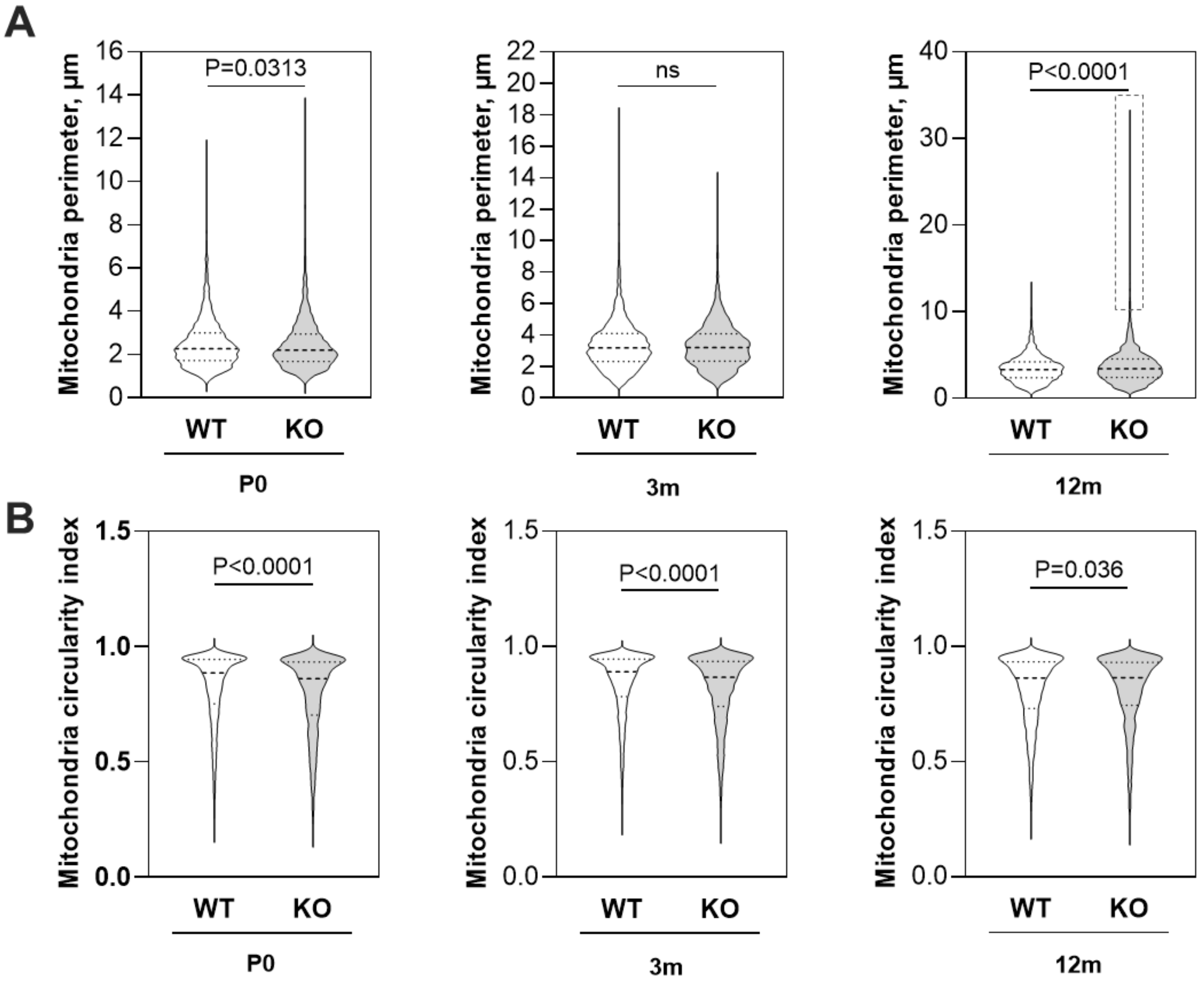


**Supplementary Figure S8.** Evaluation of the perimeter (**A**) and mitochondria circularity index (**B**) of individual mitochondria excluding fraction of damaged mitochondria. Mitochondria siting at the image border were not included in the analysis. Results are shown as violin plots, bold horizontal dashed line represents median. Statistical significance was analyzed with parametric or non-parametric t test, related to WT. Measurements in **A** and **B** were carried out on 10 images each from 3 different individuals (30 images in total). ns – not statistically significant difference. *Dashed square* in upper right violin plot indicates a population of enlarged mitochondria.

### Identification of potential MVI-interaction partners in the mouse heart

**Supplementary Table S6.** List of potential MVI-interaction proteins involved in mitochondria organization and functioning.

| No | Protein | Gene | Scores | #Peptides | References |
|---|---|---|---|---|---|
| 1 | AarF domain-containing protein kinase 1 | *Adck1* | 55.5 | 2 | *https://www.uniprot.org/uniprotkb/Q9D0L4/entry* *https://www.ncbi.nlm.nih.gov/gene/72113* |
| 2 | Reticulon-4-interacting protein 1, mitochondrial | *Rtn4ip1* | 64.8 | 2 | *https://www.uniprot.org/uniprotkb/Q924D0/entry* *https://www.ncbi.nlm.nih.gov/gene/170728* |
| 3 | Succinate dehydrogenase assembly factor 1 (complex II OXPHOS), mitochondrial | *Sdhaf1* | 57.8 | 2 | *https://www.uniprot.org/uniprotkb/Q3U276/entry* |

**Supplementary Table S7.** List of potential MVI-interaction proteins involved in autophagy.

| No | Protein | Gene | Scores | #Peptides | References |
|---|---|---|---|---|---|
| 1 | Vacuolar protein sorting-associated protein 11 homolog (Fragment) | *Vps11* | 101.2 | 2 | *https://www.uniprot.org/uniprotkb/Q91W86/entry* *https://www.ncbi.nlm.nih.gov/gene/71732* |
| 2 | WD repeat and FYVE domain-containing protein 3 | *Wdfy3* | 66.3 | 2 | *https://www.uniprot.org/uniprotkb/Q6VNB8/entry* *https://www.ncbi.nlm.nih.gov/gene/72145* |

**Supplementary Table S8.** List of potential MVI-interaction proteins involved in cellular respiration.

| No | Protein | Gene | Scores | #Peptides | References |
|---|---|---|---|---|---|
| 1 | L-lactate dehydrogenase (Fragment) | *Ldhb* | 103.3 | 2 | *https://www.uniprot.org/uniprotkb/P16125/entry* *https://www.ncbi.nlm.nih.gov/gene/16832* |
| 2 | ATP-dependent 6-phosphofructokinase, muscle type (Fragment) | *Pfkm* | 89.2 | 2 | *https://www.uniprot.org/uniprotkb/P47857/entry* *https://www.ncbi.nlm.nih.gov/gene/18642* |

**Supplementary Table S9.** List of potential MVI-interaction proteins involved in vesicle-mediated transport.

| No | Protein | Gene | Scores | #Peptides | References |
|---|---|---|---|---|---|
| 1 | AP2-associated protein kinase 1 | *Aak1* | 83.7 | 2 | *https://www.uniprot.org/uniprotkb/Q3UHJ0/entry* *https://www.ncbi.nlm.nih.gov/gene/269774* |
| 2 | BTB/POZ domain-containing protein 8 (Fragment) | *Btbd8* | 72.5 | 2 | *https://www.uniprot.org/uniprotkb/D3YUB6/entry* *https://www.ncbi.nlm.nih.gov/gene/100503185* |
| 3 | Rho GTPase-activating protein 21 | *Arhgap21* | 50.1 | 2 | *https://www.uniprot.org/uniprotkb/Q6DFV3/entry* *https://www.ncbi.nlm.nih.gov/gene/71435* |
| 4 | Stromal membrane-associated protein 2 | *Smap2* | 57.7 | 2 | *https://www.uniprot.org/uniprotkb/Q7TN29/entry* *https://www.ncbi.nlm.nih.gov/gene/69780* |
| 5 | KxDL motif-containing protein 1 (Fragment) | *Kxd1* | 84.1 | 2 | *https://www.uniprot.org/uniprotkb/Q80XH1/entry* *https://www.ncbi.nlm.nih.gov/gene/75620* |
| 6 | Vacuolar protein sorting-associated protein 51 homolog | *Vps51* | 117.5 | 2 | *https://www.uniprot.org/uniprotkb/Q3UVL4/entry* *https://www.ncbi.nlm.nih.gov/gene/68505* |
| 7 | EH domain-binding protein 1-like protein 1 (Fragment) | *Ehbp1l1* | 63.7 | 2 | *https://www.uniprot.org/uniprotkb/Q99MS7/entry* *https://www.ncbi.nlm.nih.gov/gene/114601* |
| 8 | Sorting nexin-30 | *Snx30* | 59.0 | 2 | *https://www.uniprot.org/uniprotkb/Q8CE50/entry* *https://www.ncbi.nlm.nih.gov/gene/209131* |
| 9 | TBC1 domain family member 10B | *Tbc1d10b* | 52.7 | 2 | *https://www.uniprot.org/uniprotkb/Q4KMP7/entry* *https://www.ncbi.nlm.nih.gov/gene/68449* |

| | | | | | |
|---|---|---|---|---|---|
| 10 | Brefeldin A-inhibited guanine nucleotide-exchange protein 2 | *Arfgef2* | 59.2 | 2 | *https://www.uniprot.org/uniprotkb/A2A5R2/entry* *https://www.ncbi.nlm.nih.gov/gene/99371* |
| 11 | Regulating synaptic membrane exocytosis protein 2 | *Rims2* | 61.2 | 2 | *https://www.uniprot.org/uniprotkb/Q9EQZ7/entry* *https://www.ncbi.nlm.nih.gov/gene/116838* |

**Supplementary Table S10.** List of potential MVI-interaction proteins involved in translation and protein folding.

| No | Protein | Gene | Scores | #Peptides | References |
|---|---|---|---|---|---|
| 1 | 28S ribosomal protein S15, mitochondrial | *Mrps15* | 70.9 | 2 | *https://www.ncbi.nlm.nih.gov/gene/66407* |
| 2 | Elongation factor 2 | *Eef2* | 51.1 | 2 | *https://www.uniprot.org/uniprotkb/P58252/entry* |
| 3 | RNA-splicing ligase RtcB homolog | *Rtcb* | 54.0 | 2 | *https://www.uniprot.org/uniprotkb/Q99LF4/entry* |
| 4 | Translation initiation factor eIF-2B subunit delta (Fragment) | *Eif2b4* | 66.0 | 2 | *https://www.uniprot.org/uniprotkb/Q61749/entry* |
| 5 | tRNA (cytosine(72)-C(5))-methyltransferase NSUN6 | *Nsun6* | 89.0 | 3 | *https://www.uniprot.org/uniprotkb/Q7TS68/entry* *https://www.ncbi.nlm.nih.gov/gene/74455* |
| 6 | Large subunit GTPase 1 homolog | *Lsg1* | 79.0 | 3 | *https://www.uniprot.org/uniprotkb/Q3UM18/entry* *https://www.ncbi.nlm.nih.gov/gene/224092* |
| 7 | Glutaminyl-tRNA synthetase (Fragment) | *Qars* | 50.6 | 2 | *https://www.uniprot.org/uniprotkb/Q8BML9/entry* *https://www.ncbi.nlm.nih.gov/gene/97541* |
| 8 | Kelch-like protein 20 | *Klhl20* | 82.7 | 2 | *https://www.uniprot.org/uniprotkb/Q8VCK5/entry* *https://www.ncbi.nlm.nih.gov/gene/226541* |
| 9 | Endoplasmic reticulum transmembrane helix translocase | *Atp13a1* | 57.8 | 2 | *https://www.uniprot.org/uniprotkb/Q9EPE9/entry* *https://www.ncbi.nlm.nih.gov/gene/170759* |
| 10 | Guanine nucleotide exchange factor subunit RIC1 | *Ric1* | 50.7 | 2 | *https://www.uniprot.org/uniprotkb/Q69ZJ7/entry* *https://www.ncbi.nlm.nih.gov/gene/226089* |
| 11 | Serine/threonine-protein kinase LMTK2 | *Lmtk2* | 67.9 | 2 | *https://www.uniprot.org/uniprotkb/Q3TYD6/entry* *https://www.ncbi.nlm.nih.gov/gene/231876* |

**Supplementary Table S11.** List of potential MVI-interaction proteins involved in ion transport.

| No | Protein | Gene | Scores | #Peptides | References |
|---|---|---|---|---|---|
| 1 | Aspartyl/asparaginyl beta-hydroxylase | *Asph* | 99.7 | 2 | *https://www.spandidos-publications.com/10.3892/or.2020.7792* *https://www.ncbi.nlm.nih.gov/gene?Db=gene&Cmd=ShowDetailView&TermToSearch=444* |
| 2 | Calcium channel, voltage-dependent, T type, alpha 1G subunit | *Cacna1g* | 88.5 | 2 | *https://www.uniprot.org/uniprotkb/E9PWL1/entry* *https://www.ncbi.nlm.nih.gov/gene/12291* |
| 3 | Calcium-transporting ATPase type 2C member 1 | *Atp2c1* | 74.2 | 2 | *https://www.uniprot.org/uniprotkb/Q80XR2/entry* *https://www.ncbi.nlm.nih.gov/gene/235574* |
| 4 | Canalicular multispecific organic anion transporter 2 | *Abcc3* | 84.3 | 4 | *https://www.uniprot.org/uniprotkb/B2RX12/entry* *https://pubmed.ncbi.nlm.nih.gov/10024515/* |
| 5 | H(+)/Cl(-) exchange transporter 3 | *Clcn3* | 63.6 | 2 | *https://www.uniprot.org/uniprotkb/P51791/entry* *https://www.ncbi.nlm.nih.gov/gene/12725* |
| 6 | Hyperpolarization-activated, cyclic nucleotide-gated K+ 4 | *Hcn4* | 50.7 | 3 | *https://www.uniprot.org/uniprotkb/O70507/entry* *https://www.ncbi.nlm.nih.gov/gene/330953* |
| 7 | Metalloreductase STEAP3 (Fragment) | *Steap3* | 67.2 | 2 | *https://www.uniprot.org/uniprotkb/Q8CI59/entry* *https://www.ncbi.nlm.nih.gov/gene/68428* |
| 8 | Sodium channel protein type 2 subunit alpha (Fragment) | *Scn2a* | 56.1 | 2 | *https://www.uniprot.org/uniprotkb/B1AWN6/entry* *https://www.ncbi.nlm.nih.gov/gene/110876* |

| | | | | | |
|---|---|---|---|---|---|
| 9 | Voltage-dependent calcium channel gamma-8 subunit | *Cacng8* | 87.4 | 2 | *https://www.uniprot.org/uniprotkb/Q8VHW2/entry* *https://www.ncbi.nlm.nih.gov/gene/81905* |
| 10 | Gamma-aminobutyric acid receptor subunit alpha-1 (Fragment) | *Gabra1* | 51.7 | 2 | *https://www.uniprot.org/uniprotkb/P62812/entry* *https://europepmc.org/article/med/17990949* |
| 11 | BK channel (Fragment) | *Kcnma1* | 73.5 | 2 | *https://www.uniprot.org/uniprotkb/Q08460/entry* *https://www.ncbi.nlm.nih.gov/gene/16531* |
| 12 | NHS-like protein 2 | *Nhsl2* | 53.5 | 3 | *https://www.genecards.org/cgi-bin/carddisp.pl?gene=NHSL2* |
| 13 | SH3 and cysteine-rich domain-containing protein | *Stac* | 84.8 | 2 | *https://www.uniprot.org/uniprotkb/P97306/entry* *https://www.ncbi.nlm.nih.gov/gene/20840* |

**Supplementary Table S12.** List of potential MVI-interaction proteins involved in metabolism.

| No | Protein | Gene | Scores | #Peptides | References |
|---|---|---|---|---|---|
| 1 | (Lyso)-N-acylphosphatidylethanolamine lipase (Fragment) | *Abhd4* | 63.6 | 4 | *https://www.uniprot.org/uniprotkb/Q8VD66/entry* *https://www.ncbi.nlm.nih.gov/gene/105501* |
| 2 | 17-beta-hydroxysteroid dehydrogenase 13 | *Hsd17b13* | 61.5 | 3 | *https://www.uniprot.org/uniprotkb/Q8VCR2/entry* *https://www.ncbi.nlm.nih.gov/gene/243168* |
| 3 | 1-phosphatidylinositol 4,5-bisphosphate phosphodiesterase beta-2 | *Plcb2* | 98.7 | 2 | *https://www.uniprot.org/uniprotkb/A3KGF7/entry* *https://www.ncbi.nlm.nih.gov/gene/18796* |
| 4 | Acyl-coenzyme A synthetase ACSM5, mitochondrial | *Acsm5* | 55.9 | 2 | *https://www.uniprot.org/uniprotkb/Q8BGA8/entry* *https://www.ncbi.nlm.nih.gov/gene/272428* |
| 5 | Alkyldihydroxyacetonephosphate synthase, peroxisomal | *Agps* | 68.3 | 2 | *https://www.uniprot.org/uniprotkb/Q8C0I1/entry* *https://www.ncbi.nlm.nih.gov/gene/228061* |
| 6 | Apolipoprotein L 7a | *Apol7a* | 64.6 | 2 | *https://www.ncbi.nlm.nih.gov/gene/75761* |
| 7 | Beta, beta-carotene 15,15'-dioxygenase | *Bco1* | 97.5 | 2 | *https://www.ncbi.nlm.nih.gov/gene/63857* |
| 8 | Vacuolar protein sorting-associated protein 13A (Fragment) | *Vps13a* | 106.7 | 2 | *https://www.uniprot.org/uniprotkb/Q5H8C4/entry* *https://www.ncbi.nlm.nih.gov/gene/271564* |
| 9 | Patatin-like phospholipase domain-containing protein 2 (Fragment) | *Pnpla2* | 60.4 | 3 | *https://www.uniprot.org/uniprotkb/Q8BJ56/entry* *https://www.ncbi.nlm.nih.gov/gene/66853* |
| 10 | Patatin-like phospholipase domain-containing protein 6 (Fragment) | *Pnpla6* | 80.9 | 2 | *https://www.uniprot.org/uniprotkb/Q3TRM4/entry* *https://www.ncbi.nlm.nih.gov/gene/50767* |
| 11 | Tumor necrosis factor alpha-induced protein 8 (Fragment) | *Tnfaip8* | 56.8 | 2 | *https://www.uniprot.org/uniprotkb/Q3TBL6/entry* |
| 12 | Perilipin-2 (Fragment) | *Plin2* | 75.9 | 2 | *https://www.uniprot.org/uniprotkb/P43883/entry* *https://www.ncbi.nlm.nih.gov/gene/11520* |
| 13 | Aldehyde oxidase 1 | *Aox1* | 61.3 | 2 | *https://www.uniprot.org/uniprotkb/O54754/entry* *https://www.ncbi.nlm.nih.gov/gene/11761* |
| 14 | Proprotein convertase subtilisin/kexin type 5 | *Pcsk5* | 95.5 | 2 | *https://www.uniprot.org/uniprotkb/Q04592/entry* *https://www.ncbi.nlm.nih.gov/gene/18552* |
| 15 | Carbohydrate sulfotransferase | *Chst10* | 64.5 | 2 | *https://www.uniprot.org/uniprotkb/Q6PGK7/entry* *https://www.ncbi.nlm.nih.gov/gene/98388* |
| 16 | Deoxycytidine kinase | *Dck* | 52.3 | 2 | *https://www.uniprot.org/uniprotkb/P43346/entry* *https://www.ncbi.nlm.nih.gov/gene/13178* |

**Supplementary Table S13.** List of potential MVI-interaction proteins involved in cytoskeleton organization.

| No | Protein | Gene | Scores | #Peptides | References |
|---|---|---|---|---|---|
| 1 | [F-actin]-monooxygenase MICAL2 (Fragment) | *Mical2* | 50.1 | 2 | *https://www.uniprot.org/uniprotkb/Q8BML1/entry*<br>*https://www.ncbi.nlm.nih.gov/gene?Db=gene&Cmd=DetailsSearch&Term=9645* |
| 2 | 160 kDa neurofilament protein | *Nefm* | 54.7 | 4 | *https://www.uniprot.org/uniprotkb/P08553/entry*<br>*https://www.ncbi.nlm.nih.gov/gene/18040* |
| 3 | Abl interactor 2 (Fragment) | *Abi2* | 87.1 | 2 | *https://www.uniprot.org/uniprotkb/P62484/entry*<br>*https://www.ncbi.nlm.nih.gov/gene/329165* |
| 4 | Actin, alpha skeletal muscle (Fragment) | *Acta1* | 64.2 | 2 | *https://www.uniprot.org/uniprotkb/P68134/entry*<br>*https://www.ncbi.nlm.nih.gov/gene/11459* |
| 5 | Actin, gamma-enteric smooth muscle (Fragment) | *Actg2* | 156.6 | 2 | *https://www.uniprot.org/uniprotkb/P63268/entry*<br>*https://www.ncbi.nlm.nih.gov/gene/11468* |
| 6 | Band 4.1-like protein 4B | *Epb41l4b* | 55.8 | 2 | *https://www.uniprot.org/uniprotkb/Q9JMC8/entry*<br>*https://www.ncbi.nlm.nih.gov/gene/54357* |
| 7 | Beta-adducin | *Add2* | 60.0 | 3 | *https://www.uniprot.org/uniprotkb/Q9QYB8/entry*<br>*https://www.ncbi.nlm.nih.gov/gene/11519* |
| 8 | Caldesmon 1 | *Cald1* | 60.7 | 2 | *https://www.ncbi.nlm.nih.gov/gene/109624* |
| 9 | CAP-Gly domain-containing linker protein 1 (Fragment) | *Clip1* | 95.3 | 4 | *https://www.uniprot.org/uniprotkb/Q922J3/entry*<br>*https://www.ncbi.nlm.nih.gov/gene/56430* |
| 10 | CLIP-associating protein 1 | *Clasp1* | 90.6 | 2 | *https://www.uniprot.org/uniprotkb/Q80TV8/entry*<br>*https://www.ncbi.nlm.nih.gov/gene/76707* |
| 11 | Cortactin-binding protein 2 (Fragment) | *Cttnbp2* | 75.8 | 2 | *https://www.uniprot.org/uniprotkb/B9EJA2/entry*<br>*https://www.ncbi.nlm.nih.gov/gene/30785* |
| 12 | Dynein, axonemal, heavy chain 7C | *Dnah7c* | 96.6 | 2 | *https://www.ncbi.nlm.nih.gov/gene/?term=Dnah7c* |
| 13 | Dystonin | *Dst* | 81.2 | 2 | *https://www.uniprot.org/uniprotkb/Q91ZU6/entry*<br>*https://www.ncbi.nlm.nih.gov/gene/13518* |
| 14 | F-actin monooxygenase | *Mical* | 72.2 | 3 | *https://www.uniprot.org/uniprotkb/Q8CJ19/entry*<br>*https://www.ncbi.nlm.nih.gov/gene/194401* |
| 15 | F-actin-uncapping protein LRRC16A (Fragment) | *Carmil1* | 66.5 | 2 | *https://www.uniprot.org/uniprotkb/Q6EDY6/entry*<br>*https://www.ncbi.nlm.nih.gov/gene/68732* |
| 16 | GAS2-like protein 2 | *Gas2l2* | 80.5 | 2 | *https://www.uniprot.org/uniprotkb/Q5SSG4/entry*<br>*https://www.ncbi.nlm.nih.gov/gene/237891* |
| 17 | Kinesin-like protein | *Kifc5b* | 71.4 | 2 | *https://www.uniprot.org/uniprotkb/Q9QWT9/entry*<br>*https://www.ncbi.nlm.nih.gov/gene/16580* |
| 18 | Myosin-1 | *Myh1* | 141.4 | 3 | *https://www.uniprot.org/uniprotkb/Q5SX40/entry*<br>*https://www.ncbi.nlm.nih.gov/gene/17879* |
| 19 | Nebulin | *Neb* | 69.2 | 3 | *https://www.ncbi.nlm.nih.gov/gene/17996* |
| 20 | Phosphatase and actin regulator | *Phactr2* | 59.4 | 4 | *https://www.ncbi.nlm.nih.gov/gene/215789* |
| 21 | Pleckstrin homology domain-containing, family G (with RhoGef domain) member 1 (Fragment) | *Plekhg1* | 88.5 | 2 | *https://www.uniprot.org/uniprotkb/Q4VAC9/entry*<br>*https://www.ncbi.nlm.nih.gov/gene/213783* |
| 22 | Protein ITPRID2 | *Itprid2* | 98.1 | 4 | *https://www.ncbi.nlm.nih.gov/gene/70599* |
| 23 | Protein MTSS 1 | *Mtss1* | 61.7 | 2 | *https://www.uniprot.org/uniprotkb/Q8R1S4/entry*<br>*https://www.ncbi.nlm.nih.gov/gene/211401* |
| 24 | Talin rod domain-containing protein 1 | *Tlnrd1* | 86.3 | 2 | *https://www.uniprot.org/uniprotkb/Q9ERE8/entry*<br>*https://www.ncbi.nlm.nih.gov/gene/80889* |
| 25 | Tpm1 protein | *Tpm1* | 561.2 | 9 | *https://www.uniprot.org/uniprotkb/P58771/entry*<br>*https://www.ncbi.nlm.nih.gov/gene/22003* |
| 26 | Unconventional myosin-XV | *Myo15a* | 70.7 | 3 | *https://www.uniprot.org/uniprotkb/Q9QZZ4/entry*<br>*https://www.ncbi.nlm.nih.gov/gene/17910* |

| | | | | | |
|---|---|---|---|---|---|
| 27 | Protein SOGA1 | *Soga1* | 55.5 | 3 | *https://www.uniprot.org/uniprotkb/E1U8D0/entry https://www.ncbi.nlm.nih.gov/gene/320706* |

**Supplementary Table S14.** List of potential MVI-interaction proteins involved in cell adhesion and migration.

| No | Protein | Gene | Scores | #Peptides | References |
|---|---|---|---|---|---|
| 1 | Adhesion G protein-coupled receptor L3 | *Adgrl3* | 84.1 | 2 | *https://www.uniprot.org/uniprotkb/Q80TS3/entry https://www.ncbi.nlm.nih.gov/gene/319387* |
| 2 | Carcinoembryonic antigen-related cell adhesion molecule 5 | *Ceacam5* | 98.7 | 5 | *https://www.uniprot.org/uniprotkb/Q3UKK2/entry https://www.ncbi.nlm.nih.gov/gene/73250* |
| 3 | Catenin delta-2 | *Ctnnd2* | 67.9 | 3 | *https://www.uniprot.org/uniprotkb/O35927/entry https://www.ncbi.nlm.nih.gov/gene/18163* |
| 4 | Receptor-type tyrosine-protein phosphatase alpha | *Ptpra* | 77.5 | 3 | *https://www.uniprot.org/uniprotkb/P18052/entry https://www.ncbi.nlm.nih.gov/gene/19262* |
| 5 | Receptor-type tyrosine-protein phosphatase mu | *Ptprm* | 75.6 | 2 | *https://www.uniprot.org/uniprotkb/P28828/entry https://www.ncbi.nlm.nih.gov/gene/19274* |
| 6 | Tenascin XB | *Tnxb* | 68.9 | 2 | *https://www.ncbi.nlm.nih.gov/gene/81877* |
| 7 | Vinculin | *Vcl* | 57.8 | 2 | *https://www.uniprot.org/uniprotkb/Q64727/entry https://www.ncbi.nlm.nih.gov/gene/22330* |
| 8 | Periostin (Fragment) | *Postn* | 54.3 | 2 | *https://www.uniprot.org/uniprotkb/Q62009/entry https://www.ncbi.nlm.nih.gov/gene/50706* |
| 9 | Utrophin | *Utrn* | 75.0 | 2 | *https://www.ncbi.nlm.nih.gov/gene/22288* |
| 10 | Collagen, type VI, alpha 3 | *Col6a3* | 104.4 | 3 | *https://www.ncbi.nlm.nih.gov/gene/12835* |
| 11 | Tenascin | *Tnc* | 97.3 | 2 | *https://www.uniprot.org/uniprotkb/Q80YX1/entry https://www.ncbi.nlm.nih.gov/gene/21923* |
| 12 | Tyrosine-protein kinase ABL2 | *Abl2* | 93.0 | 2 | *https://www.uniprot.org/uniprotkb/Q4JIM5/entry https://www.ncbi.nlm.nih.gov/gene/11352* |
| 13 | Symplekin | *Sympk* | 77.2 | 3 | *https://www.uniprot.org/uniprotkb/Q80X82/entry https://www.ncbi.nlm.nih.gov/gene/68188* |
| 14 | Enhancer of filamentation 1 | *Nedd9* | 51.9 | 2 | *https://www.uniprot.org/uniprotkb/O35177/entry https://www.ncbi.nlm.nih.gov/gene/18003* |
| 15 | Gliomedin | *Gldn* | 54.3 | 2 | *https://www.uniprot.org/uniprotkb/Q8BMF8/entry https://www.ncbi.nlm.nih.gov/gene/235379* |
| 16 | Protein-tyrosine-phosphatase (Fragment) | *Ptprf* | 59.4 | 2 | *https://www.uniprot.org/uniprotkb/A2A8L5/entry https://www.ncbi.nlm.nih.gov/gene/19268* |
| 17 | Filamin-A-interacting protein 1 | *Filip1* | 82.7 | 2 | *https://www.uniprot.org/uniprotkb/Q9CS72/entry https://www.ncbi.nlm.nih.gov/gene/70598* |

**Supplementary Table S15.** List of potential MVI-interaction proteins involved in cell movement.

| No | Protein | Gene | Scores | #Peptides | References |
|---|---|---|---|---|---|
| 1 | Sperm-associated antigen 17 | *Spag17* | 118.8 | 2 | *https://www.uniprot.org/uniprotkb/Q5S003/entry https://www.ncbi.nlm.nih.gov/gene/74362* |
| 2 | Bardet-Biedl syndrome 1 protein homolog | *Bbs1* | 62.1 | 2 | *https://www.uniprot.org/uniprotkb/Q3V3N7/entry https://www.ncbi.nlm.nih.gov/gene/52028* |
| 3 | Kinesin-like protein KIF27 | *Kif27* | 54.3 | 2 | *https://www.uniprot.org/uniprotkb/Q7M6Z4/entry https://www.ncbi.nlm.nih.gov/gene/75050* |
| 4 | Armadillo repeat-containing protein 3 | *Armc3* | 68.5 | 2 | *https://www.uniprot.org/uniprotkb/A2AU72/entry https://www.ncbi.nlm.nih.gov/gene/70882* |

**Supplementary Table S16.** List of potential MVI-interaction proteins involved in cell junctions.

| No | Protein | Gene | Scores | #Peptides | References |
|---|---|---|---|---|---|
| 1 | Protein HEG homolog 1 | *Heg1* | 155.0 | 2 | *https://www.uniprot.org/uniprotkb/E9Q7X6/entry* *https://www.ncbi.nlm.nih.gov/gene/77446* |
| 2 | Cingulin-like 1 | *Cgnl1* | 86.9 | 2 | *https://www.uniprot.org/uniprotkb/Q6AW69/entry* *https://www.ncbi.nlm.nih.gov/gene/68178* |
| 3 | Desmoglein-2 | *Dsg2* | 97.8 | 3 | *https://www.uniprot.org/uniprotkb/O55111/entry* *https://www.ncbi.nlm.nih.gov/gene/13511* |

**Supplementary Table S17.** List of potential MVI-interaction proteins involved in cell division.

| No | Protein | Gene | Scores | #Peptides | References |
|---|---|---|---|---|---|
| 1 | Microtubule cross-linking factor 1 (Fragment) | *Mtcl1* | 92.7 | 2 | *https://www.uniprot.org/uniprotkb/Q3UHU5/entry* *https://www.ncbi.nlm.nih.gov/gene/68617* |
| 2 | C2 domain-containing protein 3 (Fragment) | *C2cd3* | 94.2 | 2 | *https://www.uniprot.org/uniprotkb/Q52KB6/entry* *https://www.ncbi.nlm.nih.gov/gene/277939* |
| 3 | Centromere protein H | *Cenph* | 51.8 | 2 | *https://www.uniprot.org/uniprotkb/Q9QYM8/entry* *https://www.ncbi.nlm.nih.gov/gene/26886* |
| 4 | Centrosomal protein of 152 kDa | *Cep152* | 56.6 | 2 | *https://www.uniprot.org/uniprotkb/A2AUM9/entry* *https://www.ncbi.nlm.nih.gov/gene/99100* |
| 5 | Citron Rho-interacting kinase | *Cit* | 71.8 | 3 | *https://www.uniprot.org/uniprotkb/P49025/entry* *https://www.ncbi.nlm.nih.gov/gene/12704* |
| 6 | Microtubule-associated protein 10 | *Map10* | 52.0 | 2 | *https://www.uniprot.org/uniprotkb/Q8BJS7/entry* *https://www.ncbi.nlm.nih.gov/gene/74393* |
| 7 | Transforming acidic coiled-coil-containing protein 2 (Fragment) | *Tacc2* | 94.3 | 3 | *https://www.uniprot.org/uniprotkb/Q9JJG0/entry* *https://www.ncbi.nlm.nih.gov/gene/57752* |
| 8 | MutS protein homolog 4 | *Msh4* | 70.7 | 3 | *https://www.uniprot.org/uniprotkb/Q99MT2/entry* *https://www.ncbi.nlm.nih.gov/gene/55993* |
| 9 | Spindle and centriole-associated protein 1 | *Spice1* | 80.2 | 2 | *https://www.uniprot.org/uniprotkb/Q8C804/entry* *https://www.ncbi.nlm.nih.gov/gene/212514* |
| 10 | SCL-interrupting locus protein homolog | *Stil* | 82.0 | 2 | *https://www.uniprot.org/uniprotkb/Q60988/entry* *https://www.ncbi.nlm.nih.gov/gene/20460* |
| 11 | Pericentrin (Fragment) | *Pcnt* | 159.0 | 2 | *https://www.uniprot.org/uniprotkb/P48725/entry* *https://www.ncbi.nlm.nih.gov/gene/18541* |
| 12 | Probable E3 SUMO-protein ligase RNF212 | *Rnf212* | 59.7 | 2 | *https://www.uniprot.org/uniprotkb/F6TQD1/entry* *https://www.ncbi.nlm.nih.gov/gene/671564* |
| 13 | SNF-related serine/threonine-protein kinase | *Snrk* | 92.5 | 2 | *https://www.uniprot.org/uniprotkb/Q8VDU5/entry* *https://www.ncbi.nlm.nih.gov/gene/20623* |

**Supplementary Table S18.** List of potential MVI-interaction proteins involved in gene expression, transcription, and chromatin remodeling.

| No | Protein | Gene | Scores | #Peptides | References |
|---|---|---|---|---|---|
| 1 | [Histone H3]-lysine(4) N-trimethyltransferase (Fragment) | *Kmt2b* | 98.8 | 2 | *https://www.uniprot.org/uniprotkb/O08550/entry* *https://www.ncbi.nlm.nih.gov/gene/75410* |
| 2 | BarH-like 2 homeobox protein | *Barhl2* | 80.4 | 2 | *https://www.uniprot.org/uniprotkb/Q8VIB5/entry* *https://www.ncbi.nlm.nih.gov/gene/104382* |
| 3 | B-cell lymphoma/leukemia 11A | *Bcl11a* | 62.3 | 2 | *https://www.uniprot.org/uniprotkb/Q9QYE3/entry* *https://www.ncbi.nlm.nih.gov/gene/14025* |
| 4 | Bromodomain PHD finger transcription factor | *Bptf* | 77.7 | 3 | *https://www.ncbi.nlm.nih.gov/gene/207165* |

| | | | | | |
|---|---|---|---|---|---|
| 5 | Bromodomain-containing protein 1 | *Brd1* | 53.9 | 2 | *https://www.uniprot.org/uniprotkb/G5E8P1/entry* *https://www.ncbi.nlm.nih.gov/gene/223770* |
| 6 | Chromodomain-helicase-DNA-binding protein 2 | *Chd2* | 86.0 | 2 | *https://www.uniprot.org/uniprotkb/E9PZM4/entry* *https://www.ncbi.nlm.nih.gov/gene/244059* |
| 7 | CXXC-type zinc finger protein 1 | *Cxxc1* | 85.0 | 2 | *https://www.uniprot.org/uniprotkb/Q9CWW7/entry* *https://www.ncbi.nlm.nih.gov/gene/74322* |
| 8 | DNA topoisomerase 3-beta-1 | *Top3b* | 60.5 | 2 | *https://www.uniprot.org/uniprotkb/Q9Z321/entry* *https://www.ncbi.nlm.nih.gov/gene/21976* |
| 9 | Glutamine and serine-rich 1 | *Qser1* | 96.3 | 2 | *https://www.uniprot.org/uniprotkb/A0A338P6K9/entry* |
| 10 | Growth/differentiation factor 7 | *Gdf7* | 88.1 | 2 | *https://www.ncbi.nlm.nih.gov/gene/238057* |
| 11 | Histone H1.2 | *H1-2* | 52.1 | 2 | *https://www.uniprot.org/uniprotkb/P15864/entry* |
| 12 | Histone-lysine N-methyltransferase EZH2 | *Ezh2* | 65.2 | 2 | *https://www.uniprot.org/uniprotkb/Q61188/entry* *https://www.ncbi.nlm.nih.gov/gene/14056* |
| 13 | Iroquois-class homeodomain protein IRX-6 | *Irx6* | 55.5 | 2 | *https://www.uniprot.org/uniprotkb/Q9ER75/entry* *https://www.ncbi.nlm.nih.gov/gene/64379* |
| 14 | Lamina-associated polypeptide 2, isoforms beta/delta/epsilon/gamma | *Tmpo* | 67.0 | 2 | *https://www.uniprot.org/uniprotkb/Q61033/entry* |
| 15 | Menin (Fragment) | *Men1* | 52.1 | 2 | *https://www.uniprot.org/uniprotkb/O88559/entry* *https://www.ncbi.nlm.nih.gov/gene/17283* |
| 16 | NF-kappa-B-repressing factor | *Nkrf* | 58.8 | 2 | *https://www.uniprot.org/uniprotkb/Q8BY02/entry* *https://www.ncbi.nlm.nih.gov/gene/77286* |
| 17 | Nuclear receptor coactivator 6 | *Ncoa6* | 54.3 | 2 | *https://www.uniprot.org/uniprotkb/Q9JL19/entry* *https://www.ncbi.nlm.nih.gov/gene/56406* |
| 18 | Nuclear receptor corepressor 1 | *Ncor1* | 97.2 | 2 | *https://www.uniprot.org/uniprotkb/Q60974/entry* *https://www.ncbi.nlm.nih.gov/gene/20185* |
| 19 | Oligodendrocyte transcription factor 2 | *Olig2* | 103.1 | 2 | *https://www.uniprot.org/uniprotkb/Q9EQW6/entry* *https://www.ncbi.nlm.nih.gov/gene/50913* |
| 20 | Paired mesoderm homeobox protein 2 | *Prrx2* | 66.7 | 3 | *https://www.ncbi.nlm.nih.gov/gene/20204* |
| 21 | Predicted gene 6871 | *Gm6871* | 78.1 | 2 | *https://www.ncbi.nlm.nih.gov/gene/?term=Gm6871* |
| 22 | Protein CBFA2T1 | *Runx1t1* | 70.7 | 2 | *https://www.uniprot.org/uniprotkb/Q61909/entry* *https://www.ncbi.nlm.nih.gov/gene/12395* |
| 23 | Protein strawberry notch homolog 2 (Fragment) | *Sbno2* | 95.3 | 3 | *https://www.uniprot.org/uniprotkb/Q7TNB8/entry* *https://www.ncbi.nlm.nih.gov/gene/216161* |
| 24 | Regulatory factor X, 7 | *Rfx7* | 64.7 | 3 | *https://www.uniprot.org/uniprotkb/F8VPJ6/entry* *https://www.ncbi.nlm.nih.gov/gene/319758* |
| 25 | Retinoic acid-induced protein 1 | *Rai1* | 130.3 | 2 | *https://www.uniprot.org/uniprotkb/Q61818/entry* *https://www.ncbi.nlm.nih.gov/gene/19377* |
| 26 | RING finger protein 10 | *Rnf10* | 82.6 | 2 | *https://www.uniprot.org/uniprotkb/Q3UIW5/entry* *https://www.ncbi.nlm.nih.gov/gene/50849* |
| 27 | Scm polycomb group protein-like 2 | *Scml2* | 126.8 | 2 | *https://www.ncbi.nlm.nih.gov/gene/107815* |
| 28 | Set1/Ash2 histone methyltransferase complex subunit ASH2 | *Ash2l* | 95.9 | 2 | *https://www.uniprot.org/uniprotkb/Q91X20/entry* *https://www.ncbi.nlm.nih.gov/gene/23808* |
| 29 | T-complex protein 10a | *Tcp10a* | 58.7 | 2 | *https://www.ncbi.nlm.nih.gov/gene/21461* |
| 30 | Teashirt homolog 1 | *Tshz1* | 87.9 | 2 | *https://www.uniprot.org/uniprotkb/Q5DTH5/entry* *https://www.ncbi.nlm.nih.gov/gene/110796* |
| 31 | Transcription factor 12 (Fragment) | *Tcf12* | 59.4 | 2 | *https://www.uniprot.org/uniprotkb/Q61286/entry* *https://www.ncbi.nlm.nih.gov/gene/21406* |
| 32 | Transcription factor 20 | *Tcf20* | 72.3 | 2 | *https://www.uniprot.org/uniprotkb/Q9EPQ8/entry* *https://www.ncbi.nlm.nih.gov/gene/21411* |

| | | | | | |
|---|---|---|---|---|---|
| 33 | Transformation/transcription domain-associated protein | *Trrap* | 70.3 | 2 | *https://www.uniprot.org/uniprotkb/Q80YV3/entry* *https://www.ncbi.nlm.nih.gov/gene/100683* |
| 34 | Tripartite motif-containing 30C | *Trim30c* | 57.8 | 2 | *https://www.ncbi.nlm.nih.gov/gene/434219* |
| 35 | Ubinuclein-2 (Fragment) | *Ubn2* | 75.5 | 2 | *https://www.ncbi.nlm.nih.gov/gene/320538* |
| 36 | Zinc finger homeobox protein 2 | *Zfhx2* | 123.2 | 2 | *https://www.uniprot.org/uniprotkb/Q2MHN3/entry* *https://www.ncbi.nlm.nih.gov/gene/239102* |
| 37 | Zinc finger MIZ domain-containing protein 2 | *Zmiz2* | 58.3 | 3 | *https://www.uniprot.org/uniprotkb/Q8CIE2/entry* *https://www.ncbi.nlm.nih.gov/gene/52915* |
| 38 | Zinc finger protein 217 | *Zfp217* | 137.1 | 2 | *https://www.ncbi.nlm.nih.gov/gene/228913* |
| 39 | Zinc finger protein 503 | *Znf503* | 71.4 | 2 | *https://www.uniprot.org/uniprotkb/Q7TMA2/entry* |
| 40 | Zinc finger protein 516 | *Znf516* | 100.2 | 2 | *https://www.uniprot.org/uniprotkb/Q7TSH3/entry* |
| 41 | Zinc finger protein 646 (Fragment) | *Zfp646* | 92.0 | 2 | *https://www.uniprot.org/uniprotkb/Q6NV66/entry* *https://www.ncbi.nlm.nih.gov/gene/233905* |
| 42 | Zinc finger protein 827 | *Zfp827* | 65.7 | 2 | *https://www.uniprot.org/uniprotkb/Q505G8/entry* *https://www.ncbi.nlm.nih.gov/gene/622675* |
| 43 | Zinc finger protein castor homolog 1 | *Casz1* | 64.0 | 3 | *https://www.uniprot.org/uniprotkb/Q9CWL2/entry* *https://www.ncbi.nlm.nih.gov/gene/69743* |
| 44 | Pinin | *Pnn* | 69.2 | 2 | *https://www.uniprot.org/uniprotkb/O35691/entry* *https://www.ncbi.nlm.nih.gov/gene/18949* |
| 45 | LIM domain only 7 | *Lmo7* | 111.8 | 3 | *https://www.ncbi.nlm.nih.gov/gene/380928* |
| 46 | Chromodomain-helicase-DNA-binding protein 1 | *Chd1* | 87.4 | 3 | *https://www.uniprot.org/uniprotkb/P40201/entry* *https://www.ncbi.nlm.nih.gov/gene/12648* |
| 47 | Chromatin-remodeling ATPase INO80 | *Ino80* | 94.1 | 4 | *https://www.uniprot.org/uniprotkb/Q6ZPV2/entry* *https://www.ncbi.nlm.nih.gov/gene/68142* |
| 48 | Chromodomain-helicase-DNA-binding protein 8 | *Chd8* | 62.8 | 2 | *https://www.uniprot.org/uniprotkb/Q09XV5/entry* *https://www.ncbi.nlm.nih.gov/gene/67772* |
| 49 | Methylcytosine dioxygenase TET | *Tet2* | 68.8 | 2 | *https://www.uniprot.org/uniprotkb/Q4JK59/entry* *https://www.ncbi.nlm.nih.gov/gene/214133* |
| 50 | Zinc finger transcription factor Trps1 | *Trps1* | 92.7 | 2 | *https://www.uniprot.org/uniprotkb/Q925H1/entry* *https://www.ncbi.nlm.nih.gov/gene/83925* |
| 51 | Regulation of nuclear pre-mRNA domain-containing protein 2 (Fragment) | *Rprd2* | 67.0 | 3 | *https://www.ncbi.nlm.nih.gov/gene/75137* |
| 52 | Serrate RNA effector molecule homolog (Fragment) | *Srrt* | 85.7 | 2 | *https://www.uniprot.org/uniprotkb/Q99MR6/entry* *https://www.ncbi.nlm.nih.gov/gene/83701* |
| 53 | Leucine-rich repeat-containing protein 34 | *Lrrc34* | 56.9 | 2 | *https://www.uniprot.org/uniprotkb/Q9DAM1/entry* |
| 54 | ATM interactor | *Atmin* | 77.2 | 4 | *https://www.uniprot.org/uniprotkb/Q6P9S1/entry* *https://www.ncbi.nlm.nih.gov/gene/234776* |
| 55 | RuvB-like helicase (Fragment) | *Ruvbl2* | 53.4 | 2 | *https://www.uniprot.org/uniprotkb/Q9WTM5/entry* *https://www.ncbi.nlm.nih.gov/gene/20174* |

**Supplementary Table S19.** List of potential MVI-interaction proteins involved in RNA processing.

| No | Protein | Gene | Scores | #Peptides | References |
|---|---|---|---|---|---|
| 1 | PAN2-PAN3 deadenylation complex catalytic subunit Pan2 | *Pan2* | 55.5 | 2 | *https://www.uniprot.org/uniprotkb/Q8BGF7/entry* *https://www.ncbi.nlm.nih.gov/gene/103135* |
| 2 | Pre-mRNA-processing factor 17 | *Cdc40* | 77.8 | 3 | *https://www.uniprot.org/uniprotkb/Q9DC48/entry* *https://www.ncbi.nlm.nih.gov/gene/71713* |
| 3 | Splicing factor 3B subunit 1 | *Sf3b1* | 59.2 | 2 | *https://www.uniprot.org/uniprotkb/Q99NB9/entry* *https://www.ncbi.nlm.nih.gov/gene/81898* |

| | | | | | |
|---|---|---|---|---|---|
| 4 | Splicing factor 45 (Fragment) | *Rbm17* | 55.0 | 2 | *https://www.uniprot.org/uniprotkb/Q8JZX4/entry* *https://www.ncbi.nlm.nih.gov/gene/76938* |
| 5 | Tuftelin-interacting protein 11 | *Tfip11* | 58.2 | 2 | *https://www.uniprot.org/uniprotkb/Q9ERA6/entry* *https://www.ncbi.nlm.nih.gov/gene/54723* |
| 6 | Heterogeneous nuclear ribonucleoprotein A3 (Fragment) | *Hnrnpa3* | 83.5 | 2 | *https://www.uniprot.org/uniprotkb/Q8BG05/entry* *https://www.ncbi.nlm.nih.gov/gene/229279* |
| 7 | Heterogeneous nuclear ribonucleoprotein M (Fragment) | *Hnrnpm* | 108.4 | 2 | *https://www.uniprot.org/uniprotkb/Q9D0E1/entry* *https://www.ncbi.nlm.nih.gov/gene/76936* |
| 8 | RNA-binding protein 27 | *Rbm27* | 147.2 | 2 | *https://www.uniprot.org/uniprotkb/Q5SFM8/entry* *https://www.ncbi.nlm.nih.gov/gene/225432* |
| 9 | RNA-binding protein 38 | *Rbm38* | 66.2 | 2 | *https://www.uniprot.org/uniprotkb/Q62176/entry* *https://www.ncbi.nlm.nih.gov/gene/56190* |
| 10 | Integrator complex subunit 10 | *Ints10* | 78.0 | 2 | *https://www.uniprot.org/uniprotkb/Q8K2A7/entry* *https://www.ncbi.nlm.nih.gov/gene/70885* |
| 11 | Integrator complex subunit 6 | *Ints6* | 66.2 | 2 | *https://www.uniprot.org/uniprotkb/Q6PCM2/entry* *https://www.ncbi.nlm.nih.gov/gene/18130* |
| 12 | Putative RNA-binding protein 15B | *Rbm15b* | 133.3 | 2 | *https://www.uniprot.org/uniprotkb/Q6PHZ5/entry* *https://www.ncbi.nlm.nih.gov/gene/109095* |
| 13 | AHNAK nucleoprotein 2 (Fragment) | *Ahnak2* | 77.9 | 2 | *https://www.ncbi.nlm.nih.gov/gene/100041194* |
| 14 | ESF1 homolog | *Esf1* | 82.4 | 3 | *https://www.uniprot.org/uniprotkb/Q3V1V3/entry* *https://www.ncbi.nlm.nih.gov/gene/66580* |
| 15 | mRNA (2'-O-methyladenosine-N(6)-)-methyltransferase | *Pcif1* | 68.4 | 2 | *https://www.uniprot.org/uniprotkb/P59114/entry* *https://www.ncbi.nlm.nih.gov/gene/228866* |
| 16 | Nuclear pore complex protein Nup155 | *Nup155* | 74.8 | 2 | *https://www.uniprot.org/uniprotkb/Q99P88/entry* *https://www.ncbi.nlm.nih.gov/gene/170762* |
| 17 | Nucleoporin NUP188 | *Nup188* | 84.2 | 3 | *https://www.uniprot.org/uniprotkb/Q6ZQH8/entry* *https://www.ncbi.nlm.nih.gov/gene/227699* |
| 18 | Interleukin enhancer-binding factor 3 | *Ilf3* | 57.4 | 2 | *https://www.uniprot.org/uniprotkb/Q9Z1X4/entry* *https://www.ncbi.nlm.nih.gov/gene/16201* |
| 19 | TATA-box-binding protein-associated factor 15 (Fragment) | *Taf15* | 120.2 | 2 | *https://www.ncbi.nlm.nih.gov/gene/70439* |
| 20 | Schlafen family member 8 (Fragment) | *Slfn8* | 100.7 | 3 | *https://www.uniprot.org/uniprotkb/B1ARD8/entry* *https://www.ncbi.nlm.nih.gov/gene/276950* |
| 21 | Putative ATP-dependent RNA helicase DHX57 | *Dhx57* | 90.9 | 2 | *https://www.uniprot.org/uniprotkb/Q6P5D3/entry* *https://www.ncbi.nlm.nih.gov/gene/106794* |

**Supplementary Table S20.** List of potential MVI-interaction proteins involved in replication and DNA repair.

| No | Protein | Gene | Scores | #Peptides | References |
|---|---|---|---|---|---|
| 1 | Transmembrane protein 39B | *Tmem39b* | 77.5 | 2 | *https://www.uniprot.org/uniprotkb/Q810L4/entry* *https://www.ncbi.nlm.nih.gov/gene/230770* |
| 2 | Fanconi anemia group B protein homolog | *Fancb* | 63.5 | 2 | *https://www.uniprot.org/uniprotkb/Q5XJY6/entry* |
| 3 | Serine/threonine-protein kinase ATR | *Atr* | 54.1 | 2 | *https://www.uniprot.org/uniprotkb/Q9JKK8/entry* *https://www.ncbi.nlm.nih.gov/gene/245000* |
| 4 | Serine-protein kinase ATM | *Atm* | 59.1 | 2 | *https://www.uniprot.org/uniprotkb/Q62388/entry* *https://www.ncbi.nlm.nih.gov/gene/11920* |
| 5 | DNA excision repair protein ERCC-6 | *Ercc6* | 97.2 | 3 | *https://www.uniprot.org/uniprotkb/F8VPZ5/entry* *https://www.ncbi.nlm.nih.gov/gene/319955* |
| 6 | DNA ligase 4 | *Lig4* | 51.0 | 2 | *https://www.uniprot.org/uniprotkb/Q8BTF7/entry* *https://www.ncbi.nlm.nih.gov/gene/319583* |
| 7 | DNA-directed DNA/RNA polymerase mu | *Polm* | 63.8 | 2 | *https://www.uniprot.org/uniprotkb/Q9JIW4/entry* *https://www.ncbi.nlm.nih.gov/gene/54125* |

| | | | | | |
|---|---|---|---|---|---|
| 8 | Treslin | *Ticrr* | 55.8 | 4 | *https://www.uniprot.org/uniprotkb/Q8BQ33/entry*<br>*https://www.ncbi.nlm.nih.gov/gene/77011* |
| 9 | DNA polymerase subunit gamma-1 | *Polg* | 119.7 | 2 | *https://www.ncbi.nlm.nih.gov/gene/18975* |
| 10 | Chromatin assembly factor 1 subunit A | *Chaf1a* | 71.3 | 2 | *https://www.uniprot.org/uniprotkb/Q9QWF0/entry*<br>*https://www.ncbi.nlm.nih.gov/gene/27221* |
| 11 | DNA polymerase nu | *Poln* | 102.4 | 2 | *https://www.uniprot.org/uniprotkb/Q7TQ07/entry*<br>*https://www.ncbi.nlm.nih.gov/gene/272158* |
| 12 | DNA/RNA-binding protein KIN17 | *Kin* | 78.6 | 2 | *https://www.uniprot.org/uniprotkb/Q8K339/entry*<br>*https://www.ncbi.nlm.nih.gov/gene/16588* |

**Supplementary Table S21.** List of potential MVI-interaction proteins involved in stress response.

| No | Protein | Gene | Scores | #Peptides | References |
|---|---|---|---|---|---|
| 1 | Heat shock protein HSP 90-beta | *Hsp90ab1* | 50.7 | 5 | *https://www.uniprot.org/uniprotkb/P11499/entry*<br>*https://www.ncbi.nlm.nih.gov/gene/15516* |
| 2 | Clusterin-like protein 1 | *Clul1* | 57.8 | 2 | *https://www.genecards.org/cgi-bin/carddisp.pl?gene=CLUL1* |
| 3 | Ubiquitin-like modifier-activating enzyme 5 | *Uba5* | 71.1 | 2 | *https://www.uniprot.org/uniprotkb/Q8VE47/entry*<br>*https://www.ncbi.nlm.nih.gov/gene/66663* |

**Supplementary Table S22.** List of potential MVI-interaction proteins involved in apoptosis.

| No | Protein | Gene | Scores | #Peptides | References |
|---|---|---|---|---|---|
| 1 | Death domain-containing protein CRADD | *Cradd* | 50.8 | 2 | *https://www.uniprot.org/uniprotkb/O88843/entry*<br>*https://www.ncbi.nlm.nih.gov/gene/12905* |
| 2 | Caspase recruitment domain family, member 6 | *Card6* | 65.8 | 3 | *https://www.uniprot.org/uniprotkb/Q9D1X0/entry*<br>*https://www.ncbi.nlm.nih.gov/gene/239319* |
| 3 | PRKC apoptosis WT1 regulator protein | *Pawr* | 52.8 | 4 | *https://www.uniprot.org/uniprotkb/Q925B0/entry*<br>*https://www.ncbi.nlm.nih.gov/gene/114774* |
| 4 | Apoptotic protease-activating factor 1 | *Apaf1* | 61.4 | 2 | *https://www.uniprot.org/uniprotkb/O88879/entry*<br>*https://www.ncbi.nlm.nih.gov/gene/11783* |

**Supplementary Table S23.** List of potential MVI-interaction proteins involved in signal transduction.

| No | Protein | Gene | Scores | #Peptides | References |
|---|---|---|---|---|---|
| 1 | DmX-like protein 2 | *Dmxl2* | 60.5 | 2 | *https://www.uniprot.org/uniprotkb/Q8BPN8/entry*<br>*https://www.ncbi.nlm.nih.gov/gene/235380* |
| 2 | ATP-dependent RNA helicase DDX3X | *Ddx3x* | 50.3 | 2 | *https://www.uniprot.org/uniprotkb/Q62167/entry*<br>*https://www.ncbi.nlm.nih.gov/gene/13205* |
| 3 | Attractin | *Atrn* | 72.9 | 2 | *https://www.uniprot.org/uniprotkb/Q9WU60/entry*<br>*https://www.ncbi.nlm.nih.gov/gene/8455* |
| 4 | BTB/POZ domain-containing protein 10 | *Btbd10* | 82.0 | 2 | *https://www.uniprot.org/uniprotkb/Q80X66/entry*<br>*https://www.ncbi.nlm.nih.gov/gene/68815* |
| 5 | Cyclic nucleotide ras GEF | *Rapgef2* | 64.6 | 2 | *https://www.uniprot.org/uniprotkb/Q8CHG7/entry*<br>*https://www.ncbi.nlm.nih.gov/gene/76089* |
| 6 | Dual-specificity mitogen-activated protein kinase kinase 4 | *Map2k4* | 165.5 | 4 | *https://www.uniprot.org/uniprotkb/P47809/entry*<br>*https://www.ncbi.nlm.nih.gov/gene/26398* |
| 7 | Mitogen-activated protein kinase kinase kinase 11 | *Map3k11* | 102.2 | 2 | *https://www.uniprot.org/uniprotkb/Q80XI6/entry*<br>*https://www.ncbi.nlm.nih.gov/gene/26403* |
| 8 | Phosphatidylinositol 4-phosphate 5-kinase type-1 alpha | *Pip5k1a* | 51.0 | 2 | *https://www.uniprot.org/uniprotkb/P70182/entry*<br>*https://www.ncbi.nlm.nih.gov/gene/18720* |

| | | | | | |
|---|---|---|---|---|---|
| 9 | Protein NLRC3 (Fragment) | *Nlrc3* | 58.7 | 3 | *https://www.uniprot.org/uniprotkb/Q5DU56/entry* *https://www.ncbi.nlm.nih.gov/gene/268857* |
| 10 | Ras-GEF domain-containing family member 1B | *Rasgef1b* | 57.3 | 2 | *https://www.uniprot.org/uniprotkb/Q8JZL7/entry* *https://www.ncbi.nlm.nih.gov/gene/320292* |
| 11 | SH2B adapter protein 2 | *Sh2b2* | 63.3 | 2 | *https://www.uniprot.org/uniprotkb/Q9JID9/entry* *https://www.ncbi.nlm.nih.gov/gene/23921* |
| 12 | Tectonic-1 | *Tctn1* | 62.1 | 2 | *https://www.uniprot.org/uniprotkb/Q8BZ64/entry* *https://www.ncbi.nlm.nih.gov/gene/654470* |
| 13 | Neuronal tyrosine-phosphorylated phosphoinositide-3-kinase adapter 2 | *Nyap2* | 79.6 | 3 | *https://www.uniprot.org/uniprotkb/Q8BM65/entry* *https://www.ncbi.nlm.nih.gov/gene/241134* |
| 14 | Sorbin and SH3 domain-containing protein 1 (Fragment) | *Sorbs1* | 84.1 | 2 | *https://www.uniprot.org/uniprotkb/Q62417/entry* *https://www.ncbi.nlm.nih.gov/gene/20411* |
| 15 | SRC kinase-signaling inhibitor 1 | *Srcin1* | 120.1 | 2 | *https://www.uniprot.org/uniprotkb/Q9QWI6/entry* *https://www.ncbi.nlm.nih.gov/gene/56013* |
| 16 | Nitric oxide synthase, brain | *Nos1* | 67.1 | 3 | *https://www.uniprot.org/uniprotkb/Q9Z0J4/entry* *https://www.ncbi.nlm.nih.gov/gene/18125* |
| 17 | Alpha-protein kinase 1 | *Alpk1* | 50.4 | 4 | *https://www.uniprot.org/uniprotkb/Q9CXB8/entry* *https://www.ncbi.nlm.nih.gov/gene/71481* |
| 18 | CXXC-type zinc finger protein 4 | *Cxxc4* | 91.9 | 2 | *https://www.uniprot.org/uniprotkb/Q6NXI8/entry* *https://www.ncbi.nlm.nih.gov/gene/319478* |
| 19 | Interleukin-1 receptor-associated kinase-like 2 | *Irak2* | 97.8 | 3 | *https://www.uniprot.org/uniprotkb/Q8CFA1/entry* *https://www.ncbi.nlm.nih.gov/gene/108960* |
| 20 | Low-density lipoprotein receptor-related protein 5 | *Lrp5* | 75.3 | 2 | *https://www.uniprot.org/uniprotkb/Q91VN0/entry* *https://www.ncbi.nlm.nih.gov/gene/16973* |
| 21 | Low-density lipoprotein receptor-related protein | *Lrp6* | 95.3 | 2 | *https://www.uniprot.org/uniprotkb/O88572/entry* *https://www.ncbi.nlm.nih.gov/gene/16974* |
| 22 | Nucleoredoxin | *Nxn* | 53.8 | 2 | *https://www.uniprot.org/uniprotkb/P97346/entry* *https://www.ncbi.nlm.nih.gov/gene/18230* |
| 23 | Paralemmin A kinase anchor protein | *Pakap* | 53.8 | 2 | *https://www.uniprot.org/uniprotkb/O54931/entry* |
| 24 | Protein Wnt-5a | *Wnt5a* | 60.5 | 2 | *https://www.uniprot.org/uniprotkb/P22725/entry* *https://www.ncbi.nlm.nih.gov/gene/22418* |
| 25 | RAS protein activator-like 2 | *Rasal2* | 113.8 | 3 | *https://www.ncbi.nlm.nih.gov/gene/226525* |
| 26 | WD repeat-containing protein 11 (Fragment) | *Wdr11* | 60.5 | 2 | *https://www.uniprot.org/uniprotkb/Q8K1X1/entry* *https://www.ncbi.nlm.nih.gov/gene/207425* |
| 27 | Dedicator of cytokinesis protein 1 | *Dock1* | 70.9 | 2 | *https://www.uniprot.org/uniprotkb/Q8BUR4/entry* *https://www.ncbi.nlm.nih.gov/gene/330662* |
| 28 | Non-specific serine/threonine protein kinase (Fragment) | *Mark2* | 84.7 | 2 | *https://www.uniprot.org/uniprotkb/Q05512/entry* *https://www.ncbi.nlm.nih.gov/gene/13728* |
| 29 | Non-specific serine/threonine protein kinase | *Atr* | 58.6 | 2 | *https://www.uniprot.org/uniprotkb/Q9JKK8/entry* *https://www.ncbi.nlm.nih.gov/gene/245000* |
| 30 | Non-specific serine/threonine protein kinase | *Mark3* | 88.0 | 2 | *https://www.uniprot.org/uniprotkb/Q03141/entry* *https://www.ncbi.nlm.nih.gov/gene/17169* |
| 31 | Non-specific serine/threonine protein kinase | *Obscn* | 129.7 | 2 | *https://www.uniprot.org/uniprotkb/A2AAJ9/entry* *https://www.ncbi.nlm.nih.gov/gene/380698* |
| 32 | Non-specific serine/threonine protein kinase | *Wnk2* | 108.9 | 2 | *https://www.uniprot.org/uniprotkb/Q3UH66/entry* *https://www.ncbi.nlm.nih.gov/gene/75607* |
| 33 | Arf-GAP with GTPase, ANK repeat and PH domain-containing protein 1 | *Agap1* | 66.3 | 2 | *https://www.uniprot.org/uniprotkb/Q8BXK8/entry* *https://www.ncbi.nlm.nih.gov/gene/347722* |
| 34 | Tyrosine-protein kinase Lyn | *Lyn* | 84.6 | 2 | *https://www.uniprot.org/uniprotkb/P25911/entry* *https://www.ncbi.nlm.nih.gov/gene/17096* |
| 35 | Protein NDRG4 (Fragment) | *Ndrg4* | 50.2 | 2 | *https://www.uniprot.org/uniprotkb/Q8BTG7/entry* *https://www.ncbi.nlm.nih.gov/gene/234593* |

| | | | | | |
|---|---|---|---|---|---|
| 36 | Dedicator of cytokinesis protein 7 | *Dock7* | 90.9 | 2 | *https://www.uniprot.org/uniprotkb/Q8R1A4/entry* *https://www.ncbi.nlm.nih.gov/gene/67299* |
| 37 | Discoidin, CUB and LCCL domain-containing protein 2 | *Dcbld2* | 52.6 | 2 | *https://www.ncbi.nlm.nih.gov/gene/73379* |
| 38 | Cardiomyopathy-associated protein 5 | *Cmya5* | 50.2 | 2 | *https://www.uniprot.org/uniprotkb/Q70KF4/entry* *https://www.ncbi.nlm.nih.gov/gene/76469* |
| 39 | Predicted gene 5155 | *Gm5155* | 95.4 | 4 | *https://www.ncbi.nlm.nih.gov/gene/?term=Gm5155* |
| 40 | Histocompatibility 2, Q region locus 4 | *H2-Q4* | 63.1 | 2 | *https://www.ncbi.nlm.nih.gov/gene/15015* |

**Supplementary Table S24.** List of potential MVI-interaction proteins involved in synaptic function.

| No | Protein | Gene | Scores | #Peptides | References |
|---|---|---|---|---|---|
| 1 | SH3 and multiple ankyrin repeat domains protein 1 | *Shank1* | 82.9 | 2 | *https://www.uniprot.org/uniprotkb/D3YZU1/entry* *https://www.ncbi.nlm.nih.gov/gene/243961* |
| 2 | Ephrin type-A receptor 5 | *Epha5* | 81.1 | 3 | *https://www.uniprot.org/uniprotkb/Q60629/entry* *https://www.ncbi.nlm.nih.gov/gene/13839* |
| 3 | Kalirin (Fragment) | *Kalrn* | 82.8 | 2 | *https://www.uniprot.org/uniprotkb/A2CG49/entry* *https://www.ncbi.nlm.nih.gov/gene/545156* |
| 4 | Leucine-rich repeat-containing protein 7 (Fragment) | *Lrrc7* | 50.7 | 3 | *https://www.uniprot.org/uniprotkb/Q80TE7/entry* *https://www.ncbi.nlm.nih.gov/gene/242274* |
| 5 | Protein kinase C | *Prkcg* | 59.2 | 2 | *https://www.uniprot.org/uniprotkb/P63318/entry* *https://www.ncbi.nlm.nih.gov/gene/18752* |
| 6 | Disks large-associated protein 1 (Fragment) | *Dlgap1* | 80.8 | 3 | *https://www.uniprot.org/uniprotkb/Q9D415/entry* *https://www.ncbi.nlm.nih.gov/gene/224997* |
| 7 | F-box/SPRY domain-containing protein 1 | *Fbxo45* | 141.4 | 5 | *https://www.uniprot.org/uniprotkb/Q8K3B1/entry* *https://www.ncbi.nlm.nih.gov/gene/268882* |

**Supplementary Table S25.** List of potential MVI-interaction proteins involved in ubiquitin conjugation pathway.

| No | Protein | Gene | Scores | #Peptides | References |
|---|---|---|---|---|---|
| 1 | E3 ubiquitin-protein ligase CBL | *Cbl* | 55.2 | 2 | *https://www.uniprot.org/uniprotkb/P22682/entry* *https://www.ncbi.nlm.nih.gov/gene/12402* |
| 2 | E3 ubiquitin-protein ligase HECW2 | *Hecw2* | 53.9 | 3 | *https://www.uniprot.org/uniprotkb/Q6I6G8/entry* *https://www.ncbi.nlm.nih.gov/gene/329152* |
| 3 | E3 ubiquitin-protein ligase parkin | *Prkn* | 53.7 | 6 | *https://www.uniprot.org/uniprotkb/Q9WVS6/entry* *https://www.ncbi.nlm.nih.gov/gene/50873* |
| 4 | E3 ubiquitin-protein ligase PDZRN3 | *Pdzrn3* | 66.2 | 2 | *https://www.uniprot.org/uniprotkb/Q69ZS0/entry* *https://www.ncbi.nlm.nih.gov/gene/55983* |
| 5 | E3 ubiquitin-protein ligase TRAF7 (Fragment) | *Traf7* | 68.7 | 2 | *https://www.uniprot.org/uniprotkb/Q922B6/entry* |
| 6 | E3 ubiquitin-protein ligase TRIM47 | *Trim47* | 59.5 | 2 | *https://www.uniprot.org/uniprotkb/Q8C0E3/entry* |
| 7 | E3 ubiquitin-protein ligase TRIM69 | *Trim69* | 54.4 | 2 | *https://www.uniprot.org/uniprotkb/Q80X56/entry* |
| 8 | E3 ubiquitin-protein ligase UBR4 (Fragment) | *Ubr4* | 79.2 | 2 | *https://www.uniprot.org/uniprotkb/A2AN08/entry* *https://www.ncbi.nlm.nih.gov/gene/69116* |
| 9 | F-box only protein 28 | *Fbxo28* | 80.1 | 2 | *https://www.uniprot.org/uniprotkb/Q8BIG4/entry* *https://www.ncbi.nlm.nih.gov/gene/67948* |
| 10 | Ubiquitin carboxyl-terminal hydrolase 2 | *Usp2* | 60.0 | 2 | *https://www.uniprot.org/uniprotkb/O88623/entry* *https://www.ncbi.nlm.nih.gov/gene/53376* |

| | | | | | |
|---|---|---|---|---|---|
| 11 | Ubiquitin carboxyl-terminal hydrolase 20 | *Usp20* | 78.9 | 3 | *https://www.uniprot.org/uniprotkb/Q8C6M1/entry* *https://www.ncbi.nlm.nih.gov/gene/74270* |

**Supplementary Table S26.** List of potential MVI-interaction proteins involved in other biological process.

| No | Protein | Gene | Scores | #Peptides | References |
|---|---|---|---|---|---|
| 1 | BTB/POZ domain-containing protein KCTD3 | *Kctd3* | 57.2 | 2 | *https://www.uniprot.org/uniprotkb/Q8BFX3/entry* *https://www.ncbi.nlm.nih.gov/gene/226823* |
| 2 | Mucin-6 | *Muc6* | 53.6 | 2 | *https://www.uniprot.org/uniprotkb/Q80T03/entry* |
| 3 | Tumor necrosis factor receptor superfamily member 8 | *Tnfrsf8* | 69.2 | 4 | *https://www.uniprot.org/uniprotkb/Q60846/entry* *https://www.ncbi.nlm.nih.gov/gene/21941* |
| 4 | Vomeronasal type-1 receptor | *Vmn1r13* | 51.6 | 2 | *https://www.ncbi.nlm.nih.gov/gene/?term=Vmn1r13* |
| 5 | Myeloperoxidase | *Mpo* | 55.5 | 2 | *https://www.uniprot.org/uniprotkb/P11247/entry* *https://www.ncbi.nlm.nih.gov/gene/17523* |
| 6 | Alpha-helical coiled-coil rod protein | *Cchcr1* | 57.5 | 2 | *https://www.uniprot.org/uniprotkb/Q8K2I2/entry* *https://www.ncbi.nlm.nih.gov/gene/240084* |
| 7 | Proteoglycan 4 | *Prg4* | 76.4 | 2 | *https://www.uniprot.org/uniprotkb/Q9JM99/entry* *https://www.ncbi.nlm.nih.gov/gene/96875* |
| 8 | Major facilitator superfamily domain-containing protein 6 | *Mfsd6* | 74.2 | 2 | *https://www.uniprot.org/uniprotkb/Q8CBH5/entry* *https://www.ncbi.nlm.nih.gov/gene/98682* |
| 9 | Dipeptidyl peptidase 2 | *Dpp7* | 50.9 | 2 | *https://www.uniprot.org/uniprotkb/Q9ET22/entry* *https://www.ncbi.nlm.nih.gov/gene/83768* |
| 10 | Ankyrin repeat and MYND domain-containing protein 1 | *Ankmy1* | 61.7 | 3 | *https://www.ncbi.nlm.nih.gov/gene/241158* |
| 11 | Pappalysin-1 | *Pappa* | 68.4 | 2 | *https://www.uniprot.org/uniprotkb/Q8R4K8/entry* *https://www.ncbi.nlm.nih.gov/gene/18491* |
| 12 | Pleckstrin homology domain-containing, family A member 5 (Fragment) | *Plekha5* | 75.2 | 2 | *https://www.ncbi.nlm.nih.gov/gene/109135* |
| 13 | PIH1 domain-containing protein 2 (Fragment) | *Pih1d2* | 52.3 | 2 | *https://www.ncbi.nlm.nih.gov/gene/72614* |
| 14 | Methenyltetrahydrofolate synthase domain-containing protein | *Mthfsd* | 87.1 | 3 | *https://www.ncbi.nlm.nih.gov/gene/234814* |
| 15 | Microsomal signal peptidase 25 kDa subunit (Fragment) | *Spcs2* | 90.2 | 2 | *https://www.uniprot.org/uniprotkb/Q9CYN2/entry* |
| 16 | Prickle-like protein 2 | *Prickle2* | 63.1 | 3 | *https://www.ncbi.nlm.nih.gov/gene/243548* |

**Supplementary Table S27.** List of potential MVI-interaction proteins involved in unknown functions.

| No | Protein | Gene | Scores | #Peptides |
|---|---|---|---|---|
| 1 | T cell receptor alpha joining 15 (Fragment) | *Traj15* | 85.1 | 3 |
| 2 | IQ domain-containing protein C | *Iqcc* | 54.0 | 2 |
| 3 | Family with sequence similarity 227, member A | *Fam227a* | 67.3 | 2 |
| 4 | Nucleolar protein 4 (Fragment) | *Nol4* | 64.0 | 2 |
| 5 | T cell receptor alpha joining 28 (Fragment) | *Traj28* | 71.5 | 2 |
| 6 | RIKEN cDNA 4930579F01 gene | *4930579F01Rik* | 80.4 | 3 |

| | | | | |
|---|---|---|---|---|
| 7 | Maestro heat-like repeat family member 4 | *Mroh4* | 85.5 | 2 |
| 8 | Mucin 5, subtypes A and C, tracheobronchial/gastric | *Muc5ac* | 60.3 | 3 |
| 9 | RIKEN cDNA D630039A03 gene | *D630039A03Rik* | 58.5 | 2 |
| 10 | Capping protein-inhibiting regulator of actin-like (Fragment) | *Cracdl* | 66.3 | 2 |
| 11 | Chondroitin sulfate proteoglycan 4B | *Cspg4b* | 86.0 | 2 |
| 12 | Coiled-coil domain-containing protein 137 | *Ccdc137* | 64.7 | 2 |
| 13 | Family with sequence similarity 104, member A | *Fam104a* | 71.3 | 3 |
| 14 | Glucocorticoid-induced transcript 1 protein | *Glcci1* | 142.7 | 2 |
| 15 | Death domain-containing 1 | *Dthd1* | 68.8 | 4 |
| 16 | Fam122c protein | *Fam122c* | 54.0 | 2 |
| 17 | Dystrotelin | *Dytn* | 60.1 | 2 |
| 18 | Crystallin beta-gamma domain-containing 1 | *Crybg1* | 76.3 | 6 |
| 19 | Overexpressed in colon carcinoma 1 protein homolog | - | 76.8 | 2 |
| 20 | Vomeronasal type-1 receptor | *Vmn1r167* | 51.4 | 2 |
| 21 | ADAMTS-like 3 | *Adamtsl3* | 59.2 | 2 |
| 22 | Coiled-coil domain-containing protein 60 | *Ccdc60* | 82.2 | 2 |
| 23 | Leucine-rich repeat and IQ domain-containing protein 1 | *Lrriq1* | 79.9 | 4 |
| 24 | Coiled-coil domain-containing 185 | *Ccdc185* | 57.0 | 4 |
